# Vascularized tumor on a microfluidic chip to study mechanisms promoting tumor neovascularization and vascular targeted therapies

**DOI:** 10.1101/2023.08.07.552309

**Authors:** Magdalena Skubal, Benedict Mc Larney, Ngan Bao Phung, Juan Carlos Desmaras, Abdul Vehab Dozic, Alessia Volpe, Anuja Ogirala, Camila Longo Machado, Jakob Djibankov, Vladimir Ponomarev, Jan Grimm

**Affiliations:** Molecular Pharmacology Program, Memorial Sloan Kettering Cancer Center, New York, NY, USA; Department of Radiology, Memorial Sloan Kettering Cancer Center, New York, NY, USA; Department of Pharmacology, Weill Cornell Medical College, New York, NY, USA; Department of Radiology, Weill Cornell Medical College, New York, NY, USA

**Keywords:** microfluidic chip, neovascularization, targeted therapies, confocal imaging, optoacoustic imaging

## Abstract

The cascade of events leading to tumor formation includes induction of a tumor supporting neovasculature as a primary hallmark of cancer. Developing vasculature is difficult to evaluate *in vivo* but can be captured using microfluidic chip technology and patient derived cells. Herein, we established an *on chip* approach to investigate the mechanisms promoting tumor vascularization and vascular targeted therapies via co-culture of metastatic renal cell carcinoma spheroids and endothelial cells in a 3D environment. Our model permitted real-time, high-resolution observation and assessment of tumor-induced angiogenesis, where endothelial cells sprout towards the tumor and mimic a vascular network. Bevacizumab, an angiogenic inhibitor, disrupted interactions between vessels and tumors, destroying the vascular network. The *on chip* approach enabled assessment of endothelial cell biology, vessel’s functionality, drug delivery, and molecular expression of PSMA. Finally, observations in the vascularized tumor *on chip* permitted direct and conclusive quantification of this therapy in weeks as opposed to months in a comparable animal model.

**Teaser:** Vascularized tumor on microfluidic chip provides opportunity to study targeted therapies and improves preclinical drug discovery.

## Introduction

Tumor neovascularization is one of the hallmarks of cancer (*1*) and constitutes an important area of cancer research that aims to understand how cancer cells foster the formation of new supportive blood vessels (*1, 2*). Understanding the biology of tumor associated vasculature is essential to improve vascular targeted therapies frequently investigated as a strategy for cancer treatment (*3*).

Cancer research was traditionally conducted using *in vitro* two-dimensional (2D) cell cultures complemented by *in vivo* animal studies. *In vitro* cultures are valuable tools, but they often fail to reproduce the complexity of cancer phenotypes *in vivo* (*4*). Patient-derived xenografts mimic human cancer and are a frequently used tool to evaluate novel therapies, however, they are laborious, and it is expensive to screen tumors from multiple patients routinely. Studies in animals rarely allow for repeated and precise evaluation of the complex tumor microenvironment (TME) elements, particularly the tumor vasculature. Consequently, whilst these conventional models enable the evaluation of highly specific questions, too often their limitations do not translate well to clinical outcomes (*5, 6*). A potential way to approach this challenge is to replicate the functional microenvironment *in vitro* using a combination of human derived cultures and microfluidic chip (MFC) technology (*7*). In particular, three-dimensional (3D) cultures such as cell spheroids, organoids and tumoroids provide relevant and highly controllable cancer models to expedite drug discovery and evaluation (*8–11*, *5*). The MFC technology may not completely replace established *in vitro* and *in vivo* methods but is a powerful supplement permitting rapid insights. The MFC technology enables the replication of the physiology and pathophysiology of individual human organs such as blood vessels (*12–14*), brain (*15–17*), heart (*18*), lung (*19, 20*), intestine (*21, 22*), liver (*23, 24*), kidney (*18, 25*), or multiple organs connected by vascular flow (*26*). Furthermore, the nature of the MFC provides an opportunity for precision medicine, such as patient specific assessment of drug response, or personalized strategies for disease prevention (*27*, *28*).

In this work, we established an approach to investigate both tumor neovascularization and therapy via co-culture of human derived endothelial cells (EC) together with adjacently localized cancer cell spheroids on a commercially available MFC system. The small footprint of the MFC technology, up to 12 replicates per incubator, no need for animal protocol approval, quick turnaround, and versatility allows research by labs unable to access animal facilities. The MFC technology enables the cutting-edge research without reducing its impact or relevance. Additionally, the commercial availability of the system used here aids in ensuring replicability and multi-lab use. In this MFC model, there are no preformed vascular structures or guides other than two primary channels that are seeded with EC and perfused continuously with culture medium, comparable to other MFC systems (*29, 30*). Cancer cell spheroids adjacent to these primary vessels, functioning as an early tumor, induce vessels to sprout neovasculature towards the tumor. Therefore, all processes are following the cascade of events happening in a living being, but under controlled conditions and constant observation. The platform used in this work supports relevant culture parameters, luminally perfused microvasculature and the ability to intravascularly deliver compounds. Tumor size, degree of vascularization, and compounds delivery can be constantly monitored using confocal microscopy. Such a functionally vascularized tumor on the MFC permits modelling of the tumor along with its vasculature, whilst constant medium flow maintains mechanical forces like those experienced by cells *in vivo.* Within this system cells are migrating, coordinating, and organizing themselves into 3D tumor-vascular entities. We can readily create a simple microenvironment, monitor real time changes in vessels formation, probe the interactions of tumor and endothelial cells, and evaluate the role of important effectors in tumor vasculature. For example, Prostate Specific Membrane Antigen (PSMA) that emerged as a marker of tumor associated neovasculature.

Renal cell carcinoma (RCC) is a highly vascularized tumor and the 7th most common cancer in men, with a high mortality rate with median survival of 13 month, and less than 10% 5-year survival rate (*31*). In the case of metastatic RCC (mRCC), which comprises 30% of all RCC cases, this survival rate ranges from 0-20% (*32*). *In vivo* mRCC experimentation is hindered by slow growing models. Due to the prevalence and complexity of preclinical models, there is an unmet need for rapid and high throughput mRCC models for the assessment of treatment efficacy and biological mechanism elucidation.

PSMA is a transmembrane glycoprotein with a glutamate carboxypeptidase (GCPII) and folate hydrolase enzyme activity (*33*). Upregulated PSMA expression is a characteristic of prostate cancer and is associated with prostate cancer progression, metastasis, and poor prognosis in patients (*34–36*). PSMA is absent on the surface of normal EC, but PSMA expression in tumor neovasculature is a common feature in a variety of cancers and their metastatic sites, including brain, breast, lung, pancreas, bladder, or renal carcinomas (*37–41*). Therefore, in addition to wild type EC, we evaluated EC that overexpress PSMA. We characterized the process of neovascularization on the MFC stimulated by enhanced culture medium, associated with mRCC, and assessed EC response to vascular targeted therapy with bevacizumab via confocal microscopy imaging.

To emphasize the potential clinical relevance of mRCC on the MFC model, we compared therapy with bevacizumab on the MFC with an *in vivo* model of the same tumor. As bevacizumab is known to directly affect tumor vasculature (*42*), we assessed the vascular changes *in vivo* via Raster Scanning Optoacoustic Mesoscopy (RSOM) (*43*), a high-resolution optoacoustic imaging technology. The ability to reliably image and quantify vasculature both on the MFC and *in vivo* enabled a direct comparison of the MFC efficacy to established methods along with its advantages and limitations. As therapy, we chose bevacizumab, a humanized anti-Vascular Endothelial Growth Factor (VEGF) monoclonal antibody that selectively binds circulating VEGF and thus inhibits its binding to cell surface receptors on blood vessels (*42*). A recent study reported that bevacizumab has also the ability to neutralize murine VEGF and to inhibit angiogenesis and lymphangiogenesis in murine models (*44*). VEGF inhibition leads to a reduction in microvascular growth of tumor blood vessels and limits the blood supply to tumor tissues (*42*). We followed the response of renal cancer xenografts from the same mRCC line as used on the MFC, to systemic bevacizumab therapy with RSOM to image blood vessels. RSOM allowed us to characterize mRCC vasculature and monitor the dynamic response to vascular targeted therapy *in vivo* non-invasively in real time. We show how the tumor on the MFC allows precise and rapid studies of mechanisms driving tumor neovascularization, time and cost-efficient testing of vascular targeted therapies and seamlessly bridges the gap between *in vitro* and *in vivo* models, providing results in a much shorter time frame.

## Results

### Vascularization on a microfluidic chip

As a first step, we modeled the microenvironment with a functional vasculature on the MFC. Chips were prepared by filling the chamber with a collagen-based extracellular matrix (ECM) that supports cells growth. Prepositioned glass fibers were removed from the chamber to form two parallel channels (ø100 μm). The presence of channels allowed for subsequent seeding of cells into the ECM in a replicable and controlled manner, and further permitted to maintain gradient of nutrients between channels, by perfusing the one of the channels with medium containing additional factors. Seeded endothelial cells (EC) attached to the channel walls and formed a tube as the primary vessel, while medium passing through the channels at the constant speed simultaneously mimicked blood flow (Fig. 1a-c, Supplementary Fig. 1). Normal EC do not express PSMA, but its expression was reported in tumor associated neovasculature (*37–40*). Here, we considered both, wild type EC lacking PSMA (EC control), and EC engineered to overexpress functional PSMA (EC PSMA) (Supplementary Fig. 2a-b). To study sprouting of EC on the MFC, control or PSMA EC were seeded in the lower channel and were perfused with a regular medium, while the opposite channel remained unseeded and void of cells for perfusion with either regular medium or medium supplemented with tumor promoting phorbol 12-myristate 13-acetate (PMA) and increased concentration of growth factors: Vascular Endothelial Growth Factor (VEGF) and Fibroblast Growth Factor (FGF) (enhanced medium) (Table 1). Perfusion with enhanced medium created a gradient of tumor promoting factors fostering dynamic communication between channels and initiated EC sprouting. As a result, both EC control and EC PSMA were activated to sprout, and the presence of PSMA protein additionally enhanced EC migration (Fig. 1d, f). We observed that EC PSMA cells formed vessel like structures more avidly and sprouted over larger distances in comparison to EC control. The difference between both phenotypes was significant on day 10 and became more prominent on day 15 (Fig. 1f). Both, EC control and EC PSMA exposed to only regular medium neither sprouted in the same manner, nor formed vascular networks (Fig. 1e, g). Based on this result, we further concluded that tumor promoting factors in enhanced culture medium were essential to trigger EC sprouting. However, combination of these factors and PSMA protein distinctly enhanced rapid EC activation and sprouting.

**Fig 1:**
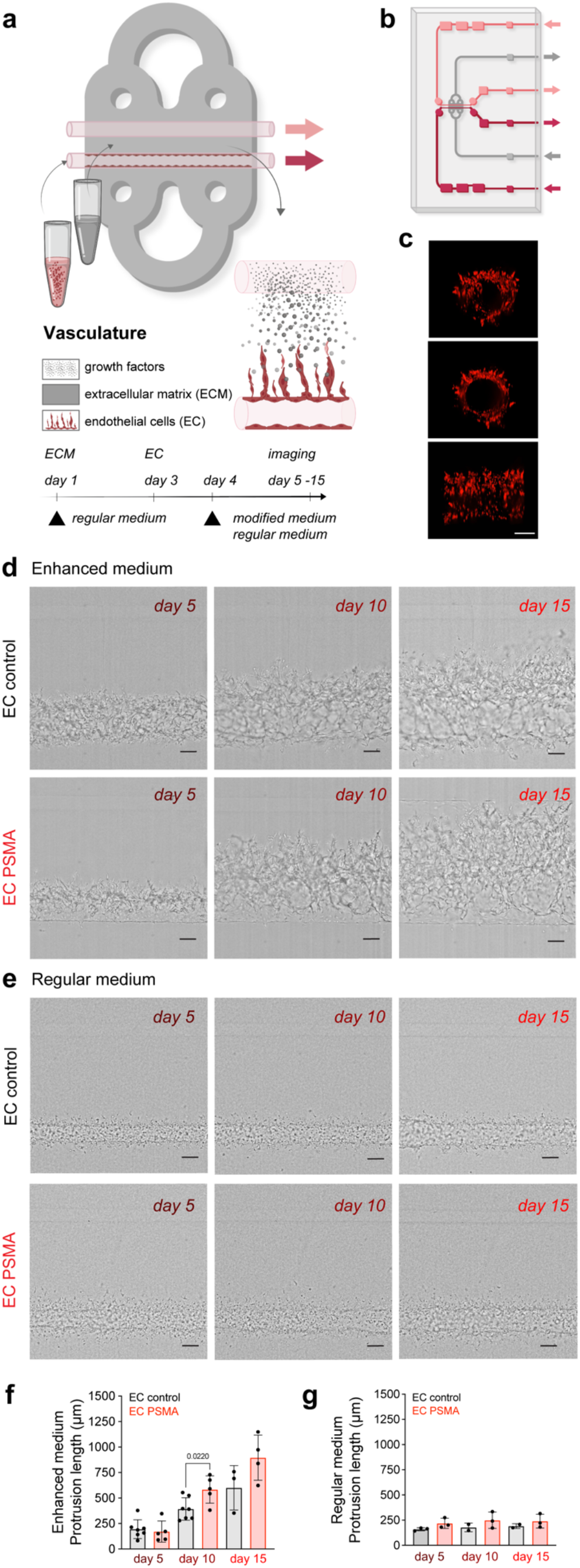
Vascularization model on the microfluidic chip. (**a**) Visualization and experimental design of vasculature on the MFC. The chip is filled with a collagen-based ECM (gray area) that supports cell growth. Two parallel channels (red and pink, ø100 μm) at the center of the chip allowed the seeding of EC and mimicked vessels. Arrows indicate the direction of constant flow (1μL/minute). (**b**) Schematic representation of the double channel MFC system built of ECM (gray area), vessels (red and pink), inlet and outlet ports (red and pink arrows) (detailed description in Supplementary Fig. 1) (**c**) Representative images of the front, side, and 3D view of the channel with red labeled EC. Scale bar 100 μm for all panels. (**d**) Confocal microscopy images showing sprouting of EC control and EC PSMA at days 5, 10 and 15. Regular medium was perfused through the lower channel seeded with EC, while enhanced medium supplemented with PMA and additional growth factors (VEGF and FGF) was perfused through the upper channel. Scale bars 100 μm. (**e**) Confocal microscopy images showing sprouting of EC control and EC PSMA cells on days 5, 10 and 15, where regular medium was perfused through both channels. Scale bars 100 μm. (**f**) Quantification of EC control (n=7) and EC PSMA (n=5) showing accelerated cell sprouting when exposed to enhanced medium. (**g**) Quantification of EC control (n=3) and EC PSMA (n=3) growth when exposed to regular medium.

**Table 1:** Composition of regular and enhanced endothelial cell medium.

| **Endothelial cell medium** | **regular** | **enhanced** |
| --- | --- | --- |
| Ascorbic Acid | 50µg/mL | 50µg/mL |
| Hydrocortisone Hemisuccinate | 1µg/mL | 1µg/mL |
| Fetal Bovine Serum (FBS) | 2% | 2% |
| L-Glutamine | 10 mM | 10 mM |
| Heparin Sulfate | 0.75 U/mL | 0.75 U/mL |
| Insulin-like Growth Factor (IGF-1) | 15 ng/mL | 15 ng/mL |
| Epidermal Growth Factor (EGF) | 5 ng/mL | 5 ng/mL |
| Fibroblast Growth Factor (<u>FGF</u>) | 5 ng/mL | 30 ng/mL |
| Vascular Endothelial Growth Factor (<u>VEGF</u>) | 5 ng/mL | 30 ng/mL |
| Phorbol 12-myristate 13-acetate (<u>PMA</u>) | - | 100ng/mL |

### Vascularized tumor on a microfluidic chip

Encouraged by the development of the sprouting vasculature on the MFC, we next established a co-culture of EC with cancer cell spheroids to address the formation of tumor associated vasculature. Expression of PSMA was identified on the surface of blood vessels associated with multiple cancers (*37, 39*), including highly vascularized renal carcinoma (*38, 45*). We chose patient-derived mRCC as a suitable tumor for the co-culture with EC since these tumors are highly vascularized. After confirming the absence of PSMA on the surface of mRCC cells (Supplementary Fig. 2c-d), we grew the tumor cells under non-adherent conditions to allow the formation of 3D cancer cell spheroids. First, mRCC spheroids mimicking early tumor stages were embedded in the ECM surrounding the chip channels, and the following day EC were seeded in both channels (Fig. 2a). To ensure optimal conditions for cell growth, the MFC was initially perfused with mRCC cell medium, and briefly before seeding the EC, the medium was changed to regular EC medium (Fig. 2a), to promote adherence of the EC and vessel formation. Co-culture of mRCC spheroids and EC on the MFC was maintained for up to 15 days (Fig. 2b). For the duration of that time mRCC spheroids were growing (Fig. 2c) and EC were sprouting towards the mRCC spheroids forming vessel like structures (Fig. 2d-e, Supplementary Fig. 3). Interestingly, independent of PSMA expression in the presence of the mRCC spheroids, EC control and EC PSMA behaved alike (Fig. 2d-e, Supplementary Fig. 4). Seeing how the MFC system modeled the vascular sprouting, we then evaluated PSMA expression on EC control. PSMA protein was absent on EC control *in vitro* and on the MFC at day 1 (Supplementary Fig. 2b, Supplementary Fig. 5), but we readily detected PSMA on EC control at day 15 on the surface of the vascular network surrounding the mRCC spheroid (Fig. 2f-j). Additional immunostainings showed an abundance of PSMA in the proximity and downstream of the mRCC spheroid indicating that factors secreted by the spheroid influenced the change of environment and induction of PSMA expression (Supplementary Fig. 6). As we characterized the model of microenvironment, we addressed the impact of flow rate on the formation of vasculature on the MFC. Cells were exposed to a constant flow of 1μL/minute, and we observed a sprouting pattern where EC growth increased with increased distance from the channel entry (highest flow, Supplementary Fig. 7).

**Fig. 2:**
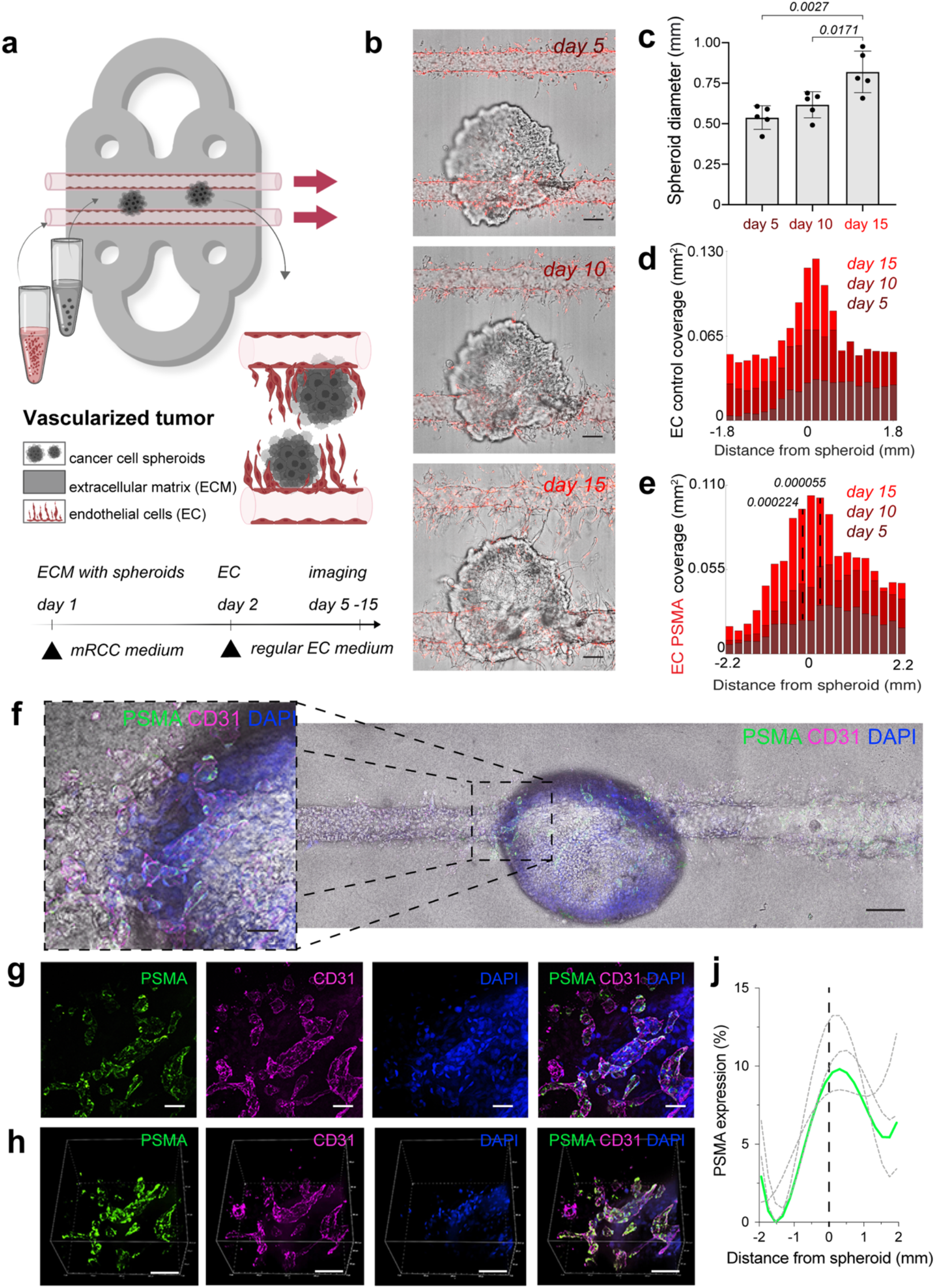
Vascularized tumor model on the microfluidic chip. (**a**) Schematic visualization and design of the vascularized tumor on the MFC. (**b**) Confocal microscopy images of EC control and mRCC spheroids co-cultured on the MFC at days 5, 10 and 15. (**c**) Quantification of mRCC spheroid growth (n=5). (**d**) Quantification of EC PSMA (n=3) and EC control (n=3) sprouting driven by mRCC spheroids. (**f**) Immunostaining showing PSMA expression on EC control cells co-cultured with mRCC spheroids for 15 days on the MFC. PSMA (green) expression was induced on the newly formed vessels (CD31, magenta) surrounding the mRCC spheroid. Endothelial and cancer cell nuclei stained with DAPI (blue). Scale bars 100 μm. (**g**) Two-dimensional and (**h**) three-dimensional visualization of PSMA (green), CD31 (magenta) and DAPI (blue) staining. Scale bars 100 μm. (**j**) Quantification of induced PSMA expression in the area surrounding mRCC spheroid (n=3).

### Assessment of vessel’s functionality and drug delivery on the microfluidic chip

To explore the functionality of tumor associated vasculature, we perfused fluorescent beads through well-developed vasculature. Beads representing an average size of typical red and white blood cells (ø10 μm and ø15 μm) were injected through the inlet port and naïvely migrated through the vascular network. Beads entered vessels penetrating the surface of the tumor spheroid, demonstrating both the vessels patency and functionality (Fig. 3a-d, Supplementary Fig. 8-9, Videos 1-3). While the beads modeled blood cells in flowing blood, we used next fluorescein to model the delivery of a small drug molecule to the tumors on the MFC. A tumor on the MFC was first perfused with fluorescein in medium for 120 min for the delivery phase, followed by perfusion with regular EC medium (for 420 min) to wash out fluorescein. Obtained time lapse images demonstrated the clear retention of fluorescence signal in the area occupied by the mRCC spheroid (Fig. 3e-g, Video 4-5).

**Fig. 3:**
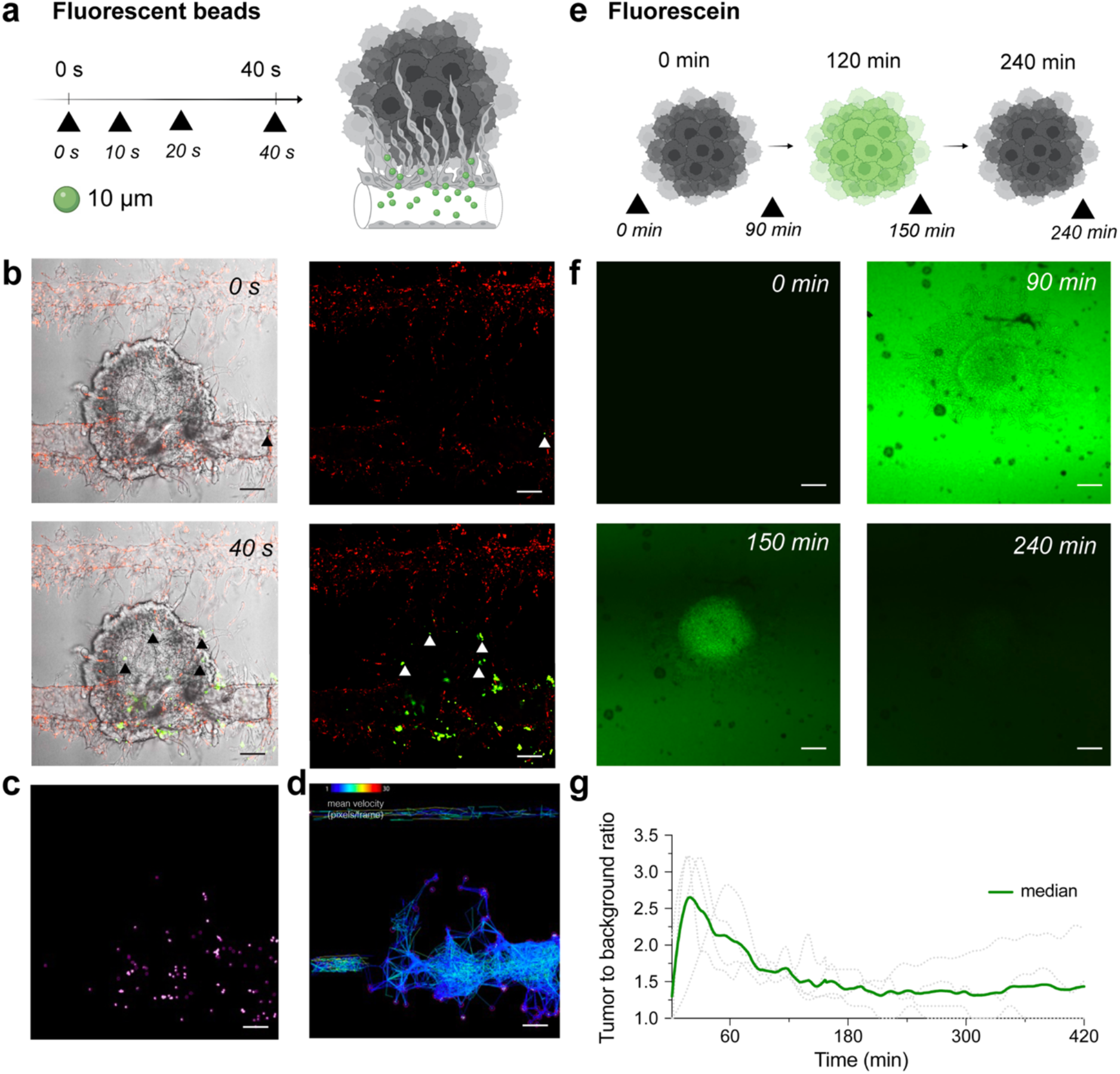
Modelling blood flow and drug delivery. (**a**) The functionality of tumor associated blood vessels was tested by perfusion of ø10 μm fluorescent beads representing the size of immune cells. Beads injected into the chip (n=1) with well-developed vasculature (day 15) through the port, entered the chips channels and newly formed vessels surrounding the surface of tumor spheroid. (**b**) Confocal microscopy images of beads traveling through the chip captured at 0s and 40s after injection (additional timepoints in Supplementary Fig. 8). Left panels represent mRCC spheroids with EC and green, fluorescent beads, while the right panels represent red labeled EC with green labeled fluorescent beads. Scale bars 100 μm. (**c**) Beads were tracked while traveling through chip channels and vasculature and became lodged within the vascular network surrounding the cancer spheroid. (**d**) Beads tracks and their mean speed. (**e**) Schematic illustration of fluorescence signal retention tested by perfusion of fluorescein (0.001 g/mL) through multiple chips (n=4) at days ranging from 5-10. Fluorescein was pumped into the chip through the bottom channel for 120 minutes followed by a washing out phase by pumping regular (fluorescein free) medium for 420 minutes. (**f**) Confocal microscopy images showing clear fluorescein retention in the mRCC spheroid at 0 minutes, 90 minutes (0-120 minutes fluorescein wash in phase), 150 minutes and 240 minutes (120-240 minutes fluorescein wash out phase). (**g**) Quantification of fluorescence signal retained in the mRCC spheroids. Scale bars 100 μm.

### Simulating vascular-targeted therapy on the microfluidic chip

The vascularized tumor on the MFC model is a promising tool for the assessment of targeted therapy as it opens the possibility to directly target endothelial cells, tumor cells, or mechanisms driving tumorigenesis that cannot be easily explored outside of *in vivo* models. Bevacizumab, a clinically approved therapeutic for advanced RCC is designed to inhibit VEGF and prevents the growth of new blood vessels. Here, mRCC spheroids and EC control cells were co-cultured on the MFC for 9 days followed by perfusion with a regular EC medium or a regular EC medium supplemented with bevacizumab (*46*). After 3 days EC in treated chips showed significant destruction, while control chips maintained in regular EC medium (bevacizumab free) throughout the duration of the experiment did not show major changes in the generated vasculature. We were able to image and reconstruct the 3D structure of the vessel highlighting the changes occurring over time (Fig. 4).

**Fig. 4:**
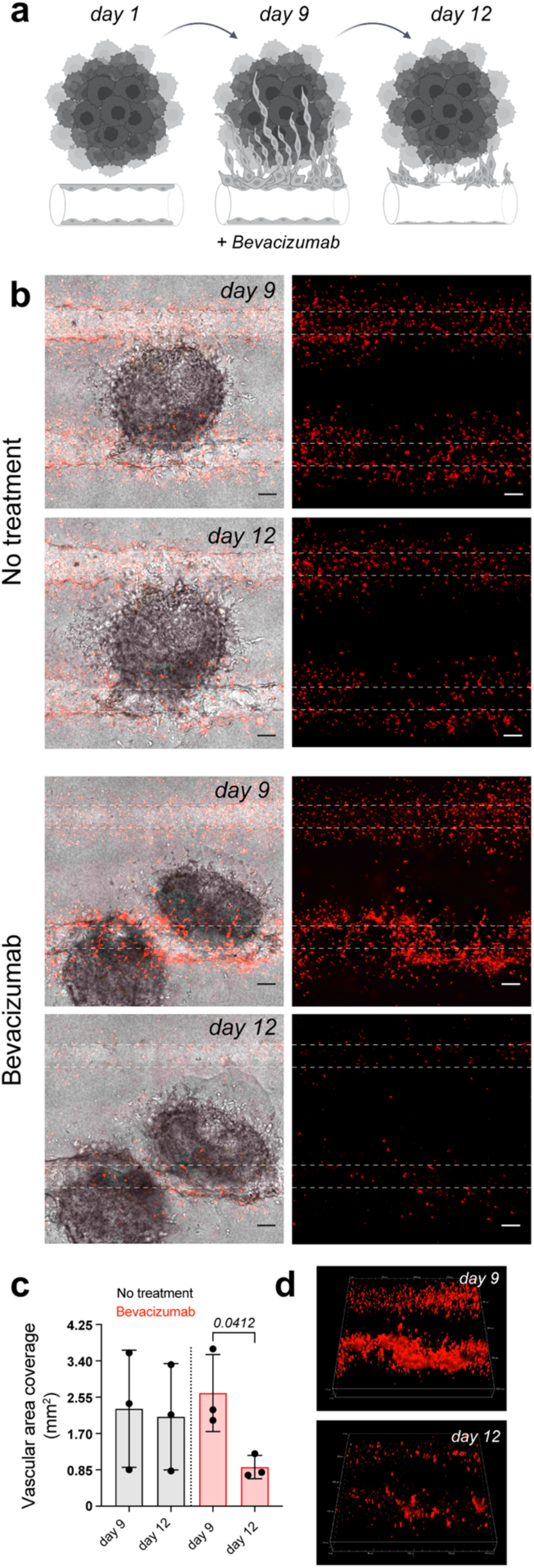
Vascular targeted therapy on the microfluidic chip. (**a**) The response to vascular targeted therapy on the MFC evaluated by treatment with bevacizumab. EC control were co-cultured with mRCC spheroids for 9 days followed by perfusion with either regular EC medium (no treatment, n=3) or EC medium supplemented with 250 μg/mL of bevacizumab (46) (n=3) for 3 days. (**b**) Confocal microscopy images of chips before (day 9) and after (day 12) bevacizumab treatment. Left panels represent mRCC spheroids (no fluorescence) with endothelial cells (red fluorescence), right panels represent endothelial cells (red fluorescence only). Scale bars 100 μm. (**c**) Quantification of cell coverage before and after treatment. (**d**) 3D visualization of EC response to bevacizumab therapy before (day 9) and after (day 12) treatment.

### Vascular targeted therapy *in vivo* as comparison

To assess the translational relevance of the tumor on the MFC approach, we compared it to an *in vivo* mouse model. Using the same mRCC line, we generated xenografts and followed the responses of mRCC tumors to bevacizumab therapy *in vivo.* Tumor volumes were monitored before (100 days) and after treatment (115 days) (Fig. 5a). We observed that bevacizumab (*47*) treated mice did not grow larger tumors in comparison to saline treated control mice (Fig. 5b). Since bevacizumab exerts a direct effect on vasculature by binding VEGF, we employed RSOM, a high-resolution optoacoustic imaging technology, to directly image and quantify blood vessels *in vivo*. RSOM enables characterizing tumor vasculature and monitoring the dynamic tumor response to targeted anti-vascular therapy (Fig. 5c, Table 2). With RSOM, we were able to assess the effectiveness of the therapy by comparing the vasculature of bevacizumab treated and control tumors, in terms of vessel length, vascular area fraction, branching, tortuosity, and nearest neighbor distance (NND) (Fig. 5d-h). We found a significant difference in vascular area fraction between treated and control tumors (Fig. 5e). By comparing the MFC system with the same tumors in an animal model, we confirmed that the MFC modeled the *in vivo* effects well and reduced the assessment of the targeted therapy from nearly 4 months *in vivo* to only a handful of days (Fig. 4a, Fig. 5a). Further *in vivo* comparison of mRCC tumor responses to bevacizumab therapy, showing reduction in vascular coverage is included in Supplementary Fig. 10.

**Fig. 5:**
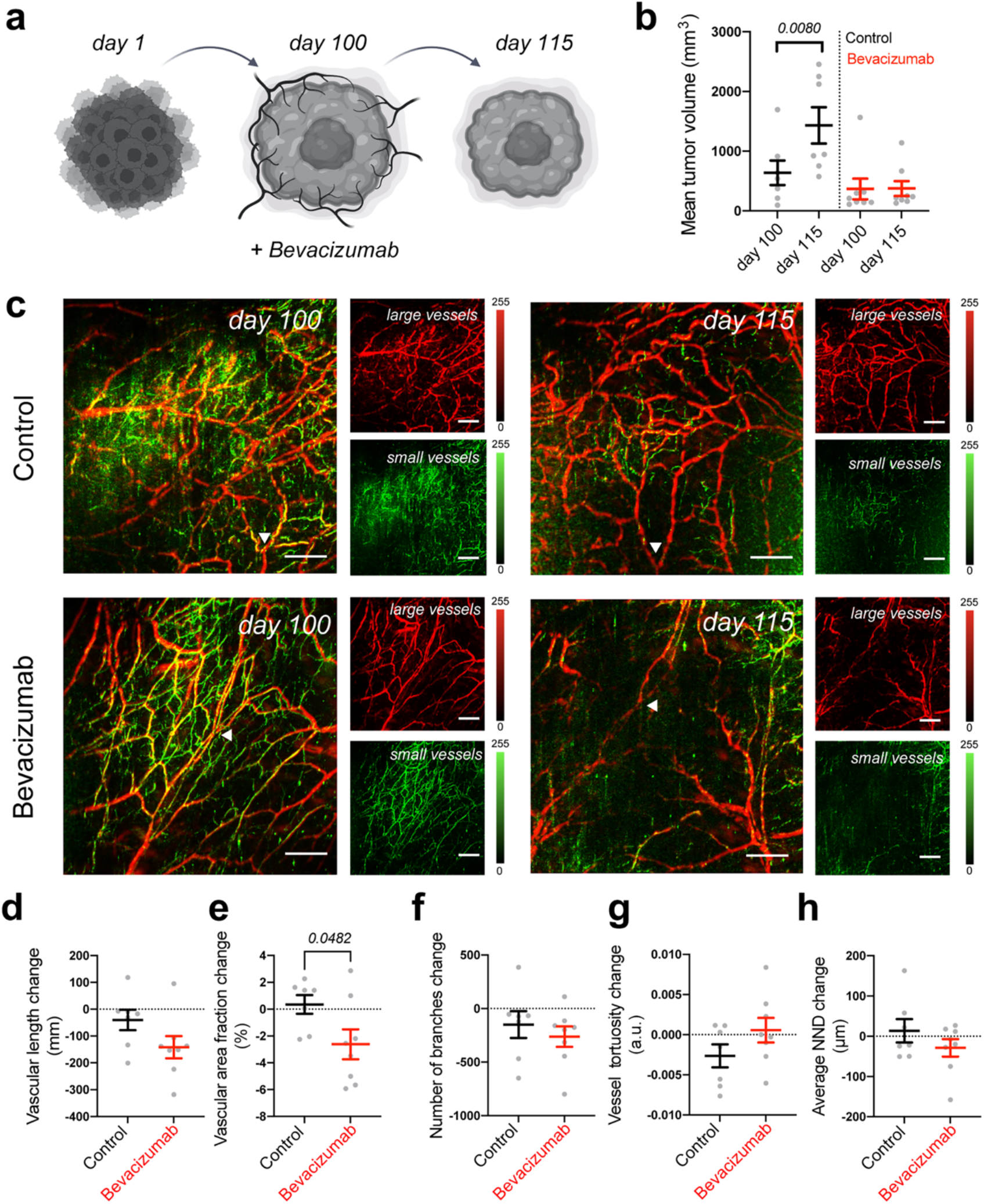
Vascular targeted therapy in vivo. (**a**) Response of mRCC mouse xenografts to vascular targeted therapy with bevacizumab. (**b**) mRCC tumors volume measured before and after treatment: mRCC tumors were grown for 100 days before receiving either saline (control group, 50uL saline each, n=7) or bevacizumab (treated group, 20 mg/kg bevacizumab (47) in saline each, n=8) 3 times a week for 2 weeks. (**c**) Representative RSOM images taken 24h before treatment commencement (day 100), and 24h after treatment completion (day 115). Images of large vessels (11-33 MHz) and small vessels (33-99 MHz). Scale bars 1mm. Quantification of (**d**) vascular length, (**e**) vascular area fraction, (**f**) number of branches, (**g**) vessel tortuosity and (**h**) nearest neighbor distance (NND).

**Table 2:**
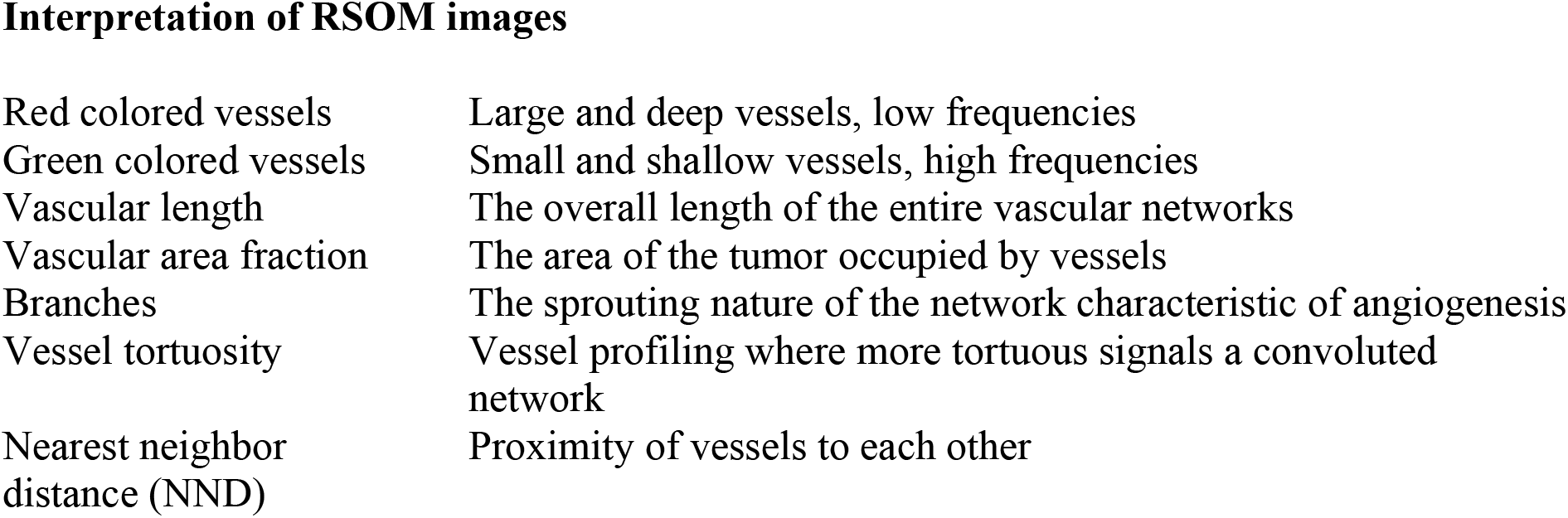
Interpretation of RSOM images.

### Assessment of angiogenesis-related proteins on the microfluidic chip

The MFC technology provides a highly controlled environment that enables facile assessment of components of the chip perfusate. Here, we employed an angiogenesis-related proteomics analysis to gain further insight into the EC biology, using the following conditions: EC control perfused with regular medium, EC control in the presence of the mRCC spheroid perfused with regular medium, EC control perfused with enriched medium, and EC PSMA perfused with enriched medium (Supplementary Fig. 11). The enriched medium contained increased levels of FGF and VEGF and accordingly, while not significant, an increasing VEGF trend was seen, with EC PSMA with enriched media having the highest expression. Also as expected, EC control and EC PSMA perfused with enriched medium had statistically higher expression levels of FGF Acidic, FGFb, FGF-4 over EC control perfused with regular medium and EC control with mRCC spheroid perfused with regular medium. However, the enriched perfused chips also had similarly increased expression of 11 proteins not added to the medium (Angiogenin, Angiopoetin-1, Activin A, ADAMTS-1, Endoglin, IGFBP-2, LAP (TGF-β1), PD-ECGF, Persephin, Prolactin, TIMP-1 and uPA). Further proteome signatures could be elucidated for all culturing conditions. We found that EC control cultured with the mRCC spheroid and perfused with regular medium had 6 proteins significantly overexpressed compared to all other conditions (DPPIV, EGF, Endothelin-1, IGFBP-3, IL-8 and Serpin E1). The EC control cultured with enriched medium proteome was also identified with significant increased expression of 5 proteins (Angiopoietin-2, Endostatin/Collagen XVIII, PDGF-AA, PDGF-AB/PDGF-BB, TIMP-4). Finally, the EC PSMA perfused with enriched medium had significantly higher expression levels of 4 proteins (EG-VEGF, IGFBP-1, MMP-9, Pentraxin 3 (PTX3)) while the EC control regular group had the highest expression for only 1 protein (GM-CSF).

## Discussion

Conventional tumor models often do not accurately reproduce the complexity that affects cancer behavior *in vivo*, and rarely allow the investigation into the role of individual elements of the TME with the visualization of how cancer cells behave when interacting with EC over time (*7*). The MFC technology bridges the gap between *in vitro* and *in vivo* research and creates a space for probing engineered microenvironments of human tissues. Here, we established an approach to model neovascularization and rapidly test vascular targeted therapies on the commercially available MFC. To emphasize its importance, we evaluated the response on the MFC to bevacizumab, a clinically approved therapeutic, and provided its direct comparison to an *in vivo* model.

Tumor EC respond to the specific TME, and cytokines mediated interactions between tumor and EC define tumor development and progression. The investigation of adaptative resistance to anti-vascular biology is essential to improve vascular targeted therapies (*48*). Normal EC do not express PSMA, but its overexpression was reported in the neovasculature of numerous cancers (*37–40*). This vascular PSMA expression urged us to consider PSMA expression in our studies. In addition to wild type EC (EC control), we included EC engineered to express PSMA (EC PSMA), mimicking tumor EC in the formation of tumor neovasculature. We generated a vascularization model on the MFC built of human EC, and a vascularized tumor model built of EC with mRCC spheroids. The vascularization on the MFC showed that PSMA positive EC responded more rapidly to environmental stimuli carried in the different culture media, resulting in fast sprouting over larger distances, however by day 15 this difference was less prominent highlighting the similarities between the EC PSMA and EC Control phenotypes (Fig. 1). Our findings are in line with previous studies reporting that PSMA positive EC foster angiogenesis by promoting tube formation, however this has so far only been shown in simplified 2D culture model (*49*). Also, our vascularization on the MFC model showed that a critical component in EC sprouting was enriched medium supplemented with additional VEGF, FGF and PMA, emphasizing the molecular investigations that can be achieved with this system. Growth factors are essential mediators in neovascularization (*50–52*), and PMA induces EC sprouting by upregulating matrix metalloproteinases, essential for cell invasion in the early stages of angiogenesis (*53*). In the vascularized tumor on the MFC, we combined EC and mRCC spheroids to create a model where endothelial and cancer cells can spontaneously migrate and self-organize to form 3D structures (Fig. 2b). Cancer spheroids allowed us to mimic aspects of *in vivo* tumor growth that are otherwise not possible to model *in vitro*, particularly neovascularization (*54, 55*). To further characterize our model, we used EC control and EC PSMA, and monitored the real time changes in vessels formation (Fig. 2d-e). The comparison between EC control and EC PSMA showed that there is no difference in the response of these cells in the presence of mRCC spheroids. EC control behaved in a similar manner to EC PSMA, aside from EC PSMA growing faster in the direct proximity of the spheroid (Supplementary Fig. 4), in line with the experimentation without spheroids.

Interestingly, the co-culture of EC control and mRCC spheroids evoked PSMA expression on EC control (Fig. 2f-j, Supplementary Fig. 6). We observed PSMA on the newly formed vessels in the proximity and downstream of the spheroid, as the factors secreted from the mRCC travel downstream of the medium flow direction on the MFC. Our studies found that co-culture of PSMA negative EC with cancer cells expressing PSMA can transform EC to express PSMA through uptake of PSMA positive membranes, such as microvesicles (*49*). Although renal carcinomas were previously used to study tumor angiogenesis on the MFC (*13*), we report for the first time that co-culture of EC and mRCC can promote PSMA expression on EC control. As PSMA expression was not observed directly on mRCC cells (Supplementary Fig. 2c-d), exposure of EC to factors present in culture medium plays an important role in microenvironment remodeling (*56*). The functional role of PSMA in angiogenesis remains elusive, but PSMA specific expression on neovasculature suggests that it participates in tumor development and progression (*57*). Even while the mechanistic details need to be explored, our data confirm that PSMA provides functional advantage to EC and aids EC sprouting (Fig. 1f). In summary, we demonstrated the potential of our MFC models to investigate neovascularization and induce neovascular PSMA expression. Furthermore, collection of perfusate permitted assessment of angiogenesis-related proteins (Supplementary Fig. 11). Importantly, comparing EC control and EC PSMA in the presence of enriched media revealed statistically higher expression levels of 11 proteins involved in angiogenesis over the other conditions. Overall, these studies showed that the presence of PSMA on the EC as well as the presence of tumors induced significant changes in the angiogenic protein signatures of the system, shifting it to a more pro-angiogenic state. Interestingly, we saw that GM-CSF decreased when control EC were co-cultured with mRCC in regular medium, and lowest when PSMA was present on the EC in enriched medium. Since GM-CSF is known to inhibit angiogenesis, this further confirms the observed sprouting results and the induction of a PSMA phenotype in EC control cells (*58*).

For personalized medicine applications, reduced throughput in exchange for higher complexity to resemble a tumor with tightly controlled intravascular delivery of candidate therapies is desirable. We first assessed vessel functionality via perfusion with fluorescent beads, representing an average size of typical red and white blood cells (Fig. 3a-d) (*59*). Beads readily traveled through channels and the vasculature surrounding mRCC spheroid, but failed to penetrate into the tumor core as often seen *in vivo* (*60*). We provided evidence that the newly formed vessels on the MFC are functional, and study of immunotherapies would be possible with this system. We next addressed the potential of the MFC to model small molecule drug delivery to tumors. To mimic non-targeted therapy, we applied the fluorescent small molecule probe, fluorescein (Fig. 3e-g) (*61*). Fluorescein is a clinically approved compound in fluorescence guided tumor resection (*62*). Fluorescein showed a clear tumor delivery, retention and wash-out phase in the MFC tumor model very similar to the mechanism seen *in vivo* with second window ICG imaging (*63, 64*), further emphasizing the recapitulation of *in vivo* environments on the MFC.

We proceeded with vascular targeted therapy. Bevacizumab is used as first line treatment in patients with mRCC where it induces a significant improvement in progression free survival (*65*). Here, we integrated *in vivo* and *in vitro* imaging methods to evaluate the response of vascularized tumor to bevacizumab, a VEGF inhibitor. Bevacizumab treatment on the MFC successfully blocked EC activation and recruitment, leaving the EC network substantially destroyed within 3 days (Fig. 4). Further *in vivo* comparison of the mRCC tumor response to bevacizumab therapy demonstrated clear differences between treated and control tumors (Fig. 5 a-b). Optoacoustic imaging enabled monitoring of *in vivo* changes in mRCC tumors vasculature (Fig. 5 c-h). After 2 weeks of treatment, we found a significant difference in vascular area fraction between treated and control tumors. In addition, vascular staining showed similar responses to bevacizumab therapy in mRCC tumors (Supplementary Fig. 10). Bevacizumab affected vessels in a comparable manner on the MFC and *in vivo* but the MFC system allowed us to grow vascularized tumors over 8 times faster than xenografts, and with higher precision to focus on mechanisms driving tumor vascularization. Thus, especially in cases of slow growing cancer xenografts like mRCC, the chip model provides a clear temporal advantage with respect to personalized medicine approaches. The MFC approach provides a rapid and reproducible method to grow human relevant models and to screen for suitable treatments, reducing experiment times from nearly 4 months to 2 weeks when compared to mouse models, a valuable time gain for cancer patients.

Results from the MFC and *in vivo* studies allowed for the performance assessment of our system and highlighted its potential for accelerated drug screening and preclinical investigation of biomarker targeting probes. Future iterations of the vascularized tumor on the MFC could facilitate investigations on tumor biology, identify ineffective therapies prior to expensive clinical trial stages, reduce the time needed for the assessment of patient xenograft models and enable a focus on patient tailored, precision cancer medicine. In addition, patient derived organoids on the MFC, used as a miniaturized tumor models and avatars for the patient’s actual tumor, would allow to recapitulate various aspects of patient physiology and disease phenotypes. These could even be combined with autologous buffy coat to add immune cells to test immunotherapies thus provide a wide opportunity for improving preclinical drug discovery, and clinical trial validation (*66, 67*). Therefore, this work has a direct impact on medical research, where it could reduce the requirement for mouse experiments and accelerate translation to clinical research if further studies confirm our findings and regulatory institution accept the impact of MFC systems. We estimate that a single dedicated incubator system could assess around 92 chips in the time required to carry out the *in vivo* experimentation in a footprint like that of two mouse cages (10 mice) increasing throughput by almost an order of magnitude. These aspects of the MFC system are particularly timely considering recent announcements by the FDA (FDA modernization act 2.0) removing the need for animal data prior to commencing clinical trials (*68*).

However, this MFC system as all preclinical models certainly has limitations. Firstly, setup does require custom installation including pumps, racks and the MFC. Secondly, the system does not contain vascular niches to model more complex tumor microenvironment, a set-up of which is possible but would require modification outside the scope of this work. Thirdly, we co-cultured only cancer and endothelial cells limiting the assessment of immune cells on this work. Additional or alternative components can be added, such as components of the immune system. Local stromal cells and pericytes were previously incorporated on this MFC (*69*). A wealth of insights and research has been possible thanks to patient-derived xenografts in immune deficient mice and MFC may readily provide further discoveries (*70*). In cases where *in vivo* insights are still required, preliminary MFC results can be used to guide and optimize this research reducing overall animal numbers used. In drug discovery pipelines the MFC could provide a reliable method for patient derived tumor growth with numerous treatments being investigated in a rapid and high throughput set-up, resulting in accelerated personalized medicine approaches for individual patients.

## Materials and Methods

### Cell culture

Primary human umbilical vein endothelial cells (EC) were purchased from Angio-proteome (#cAP-0001RFP). Endothelial cells (EC control) were engineered by retroviral gene transfer to stably overexpress functional PSMA protein (EC PSMA) (Supplementary Fig. 2). Cell culture was maintained in a humidified incubator at 37°C, 5% CO_2_ in the EC medium: VascuLife Basal Medium (Lifeline Cell Technology #LM-0002) supplemented with VEGF kit (Lifeline Cell Technology #LS-1020) containing Ascorbic Acid (50μg/mL), Hydrocortisone Hemisuccinate (1μg/mL), Fetal Bovine Serum (FBS; 2%), L-Glutamine (10 mM), Heparin Sulfate (0.75 U/mL) and human recombinant Fibroblast Growth Factor (FGF; 5 ng/mL), human recombinant Insulin-like Growth Factor (IGF-1; 15 ng/mL), human recombinant Epidermal Growth Factor (EGF; 5 ng/mL), human recombinant Vascular Endothelial Growth Factor (VEGF; 5 ng/mL). Enhanced medium was supplemented with additional VEGF (25 ng/mL), FGF (25 ng/mL) and phorbol 12-myristate 13-acetate (PMA, 100ng/mL) (Table 1). For all performed experiments EC did not exceed 3 passages.

Patient-derived metastatic renal cell carcinoma JHRCC62A (mRCC) (*71*) was kindly provided by the Abraham Hakimi Lab (MSKCC). mRCC culture was maintained in a humidified incubator at 37°C, 5% CO_2_ and was supplemented with compete DMEM/F12 medium (Corning #10-103) containing 10% Fetal Bovine Serum (GeminiBio #100-106), 1% L-Glutamine (Gibco #25030081) and 1% Penicillin/Streptomycin (Corning #30-002).

### Microfluidic system to create vascularization and vascularized tumor model

The MFC system by Nortis Inc. (Woodinville, WA) was used to establish vasculature and the vascularized tumor models. Our studies are in line with previous work with EC on this platform (*69, 72*). To model vasculature on the MFC, extracellular matrix (ECM), the foundation for the sprouting cells, was prepared by mixing chilled collagen I (Corning #354249) with phosphate buffered saline (PBS, Corning #46013CM), phenol red solution (Cepham Life Sciences #10387-0) and water to the final concentration of 7 mg/mL. Collagen was chosen on account of its mechanical stability and for its fibrillar nature. Additionally, it is less susceptible to the known inconsistencies of Matrigel and was well established for similar applications (*13, 73, 74*). Collagen density, thus pH of ECM solution was carefully adjusted to 8-8.5 by adding 1M NaOH. Prior injection of ECM to the chip, matrix chamber was washed with 100% ethanol and dried. Approximately 250μL of ECM solution was injected using chilled 1mL syringe with 20-gauge blunt needle (McMaster-Carr # 75165A677). ECM chamber was sealed, and chip was incubated overnight at room temperature to allow collagen solution to polymerize. Next day, cylindrical silica fibers were removed to form hollow channels (ø100 μm) and pre-flow (1 μL/minute) was initiated with an incubator gas pump (Nortis Inc. #IGP-001). Cells were seeded and cultured on the inner surface of the channels and within the ECM. In specific, to model the vasculature on the MFC, a single cell suspension (2μL of 10^7^cells/mL) of EC (Angio-proteome #cAP-0001RFP) was injected to the bottom channel of a double-channel MFC (Nortis Inc. #DCC-001) with a Hamilton syringe. Cells were allowed to adhere for 2h at 37°C prior to initiating flow at 1 μL/minute. Culture was maintained for up to 15 days in a humidified incubator at 37°C, 5% CO_2_. Fresh EC medium was supplied every third day.

To model the vascularized tumor on the MFC we chose mRCC cancer cell spheroids (*75*). Briefly 10,000 mRCC cells were grown in suspension on low-attachment 96 well plates (Corning #7007) to allow formations of 3D cancer cell spheroids. mRCC cells spheroids were collected 48h after seeding and gently mixed with chilled freshly prepared ECM. Next, 3D spheroids embedded in collagen-based ECM were injected into the ECM chamber surrounding two silica fibers. Chips were incubated for 3h in a humidified incubator at 37°C, 5% CO_2_. Flowing ECM polymerization, silica fibers were removed from chip to form the lumens. Pre-flow was initiated with an incubator gas pump using degassed DMEM/F12 medium (Corning #10-103). Next day DMEM/F12 medium was exchanged to EC medium (Lifeline Cell Technology #LL-0003), and EC were injected into the both lumens of a double-channel MFC. Cells were allowed to adhere for 2h at 37°C prior to initiating flow at 1 μL/minute, this sheer rate is estimated to be ∼1.05 dyn/cm^2^ within the parent vessel lumen providing physiological relevant conditions and simultaneously reducing medium consumption.

### Flow cytometry analysis

100,000 cells were washed twice with PBS and resuspended in FACS buffer (PBS pH 7.4 with 10% (v/v) FCS). Cells were stained with a human anti-PSMA monoclonal antibody in the dark (10 μg/mL, MBL #K0142-3, mouse IgG1 clone 107-1A4). Polyclonal goat anti-mouse Ig conjugated to APC was used as a secondary antibody (2 μg/mL, BD Pharmigen #550826). For all antibodies, cells were stained for 1 hour at 4 °C in FACS buffer. Flow cytometric profiles were acquired using a LSRFortessa II (BD Bioscience, USA) equipped with a FACSDiva analysis software. Data analysis was carried out using FCS Express 7 (version 7.08.0018).

### Immunocytochemical staining

30,000 cells per well were seeded in 8 well chamber glass slide (Nunc Lab-Tek II Chamber Slide System #154534). After obtaining approximately 80% of confluency cells were fixed using 4% paraformaldehyde in PBS pH 7.4 for 10 min and blocked with 10% NGS and 0.5% Triton X-100 for 1h at room temperature. Cells were washed with PBS and stained with primary mouse monoclonal anti-PSMA antibody (Abcam #187570) in the concretion 1:100 overnight at 4°C. Next day cells were washed twice with PBS and incubated with secondary anti-mouse antibodies (Alexa Fluor 488 goat anti-mouse Ig #A32723) in the concretion 1:500 for 2h. Cell nuclei were stained for 15 minutes with DAPI (Thermo Fisher Scientific #D3571). Coverslips were mounted using the PermaFluor aqueous mounting medium (Thermo Scientific # TA-006-FM).

Cells cultured on the MFC were washed with PBS and fixed with fixation/permeabilization solution (BD Biosciences #554722) for 2h at room temperature with the syringe pump (2ul/min). Next, cells were briefly rinsed with PBS and blocked with perm/wash buffer (BD Biosciences #554723) supplemented with 2% BSA and 0.05% Tween 20 for 2h at room temperature. Chips were incubated overnight at 4°C with primary antibodies: mouse monoclonal anti-PSMA (Abcam #187570) and rabbit polyclonal anti-CD31 (Abcam #32457) in the concretion 1:100 each, mixed with a fresh blocking buffer and added directly to the chip. Chips were washed with 2mL of PBS and incubated with secondary antibodies (Alexa Fluor 488 goat anti-mouse Ig #A32723, Alexa Fluor 633 goat anti-rabbit Ig #A21071) in the concretion 1:500 for 2h in room temperature. Unbound secondary antibodies were washed with PBS, cell nuclei were stained for 15 minutes with DAPI (Thermo Fisher Scientific #D3571) and chips were washed again with PBS.

### Assessment of vessels functionality and patency with fluorescent beads

To test the functionality of newly formed vessels fluorescent beads of ø10 μm (505nm/515nm) (Thermo Fisher Scientific #F8836) and ø15 μm (565nm/580nm) (Thermo Fisher Scientific #F21012) were perfused through chips at day 15. Microbeads were designed for regional blood flow determination as these matches the size of immune cells. For these types of studies beads were injected in the lower channel of the chip and traveled through the vessel-like structures and the opposite channel where the fluorescence can be tracked and quantified. Beads perfused through the chip were resistant to photobleaching and retain their fluorescence.

### Assessment of drug delivery

Fluorescein (Sigma #46955) was perfused through chips at days ranging from 5-10 to model drug delivery on the MFC. Fluorescein solution at the final concentration of 0.001g/mL in regular EC medium was washed in for 2h at 1μL/minute through the lower channel of the chip, while the upper port remined closed. Next, lower channel with fluorescein solution was closed and upper port containing only EC medium was opened to wash out fluorescein. Fluorescein was washed out from ECM after 30 minutes and retained only in the spheroid.

### Vascular targeted therapy on the microfluidic chip

Co-culture of patient derived mRCC spheroids and EC control cells was maintained until vessels were formed. 250 μg/mL of bevacizumab (*46*) was added to EC medium of treated chips, while control chips were perfused with regular EC medium. Cell culture was maintained for another 3 days. Cell coverage was quantified at the beginning of the experiment (day 9), and after 3 days of treatment with bevacizumab (day 12).

### Confocal microscopy, image generation, quantification, and analysis

Images were captured on a spinning disc confocal Nikon ECLIPSE Ti microscope equipped with EMCCD camera (Andor sCMOS) and laser (Lumencor SPECTRA Light Engine). Chips were imaged in a specially modified stage holder enabling chips to be imaged in their housing with sustained environmental conditions and pump connection. Confocal images were captured under white light conditions and 405nm for DAPI channel (excitation 401nm/emission 421nm), 488 nm for PSMA channel (excitation 496nm/emission 519nm), 561nm for RFP (excitation 555 nm/ emission 584nm), 633nm for CD31 channel (excitation 632nm/emission 647nm).

The native Nikon format was converted to 16-bit gray tiff files for quantification. In cases of suboptimal illumination conditions and to correct for stitching artefacts a fast Fourier bandpass transform was applied in Fiji (ImageJ, version 2.0.0-rc-65/1.53c). Following image correction images were imported to MATLAB (MathWorks, version 2020b) for quantification via custom scripts. Sprouting length images were firstly thresholded and cropped followed by 2D median filtering (kernel size of 20×20 pixels) to remove artefacts. The image was then dilated using a disk-shaped dilation factor. To determine the average sprouting length the region prop’s function was employed to find ovular shapes the width of the image. The circumference of this oval shape accurately traced the average projection of the sprouting cells. The distance between the outer most point of this oval circumference and the upper most channel point was calculated to give the average sprouting distance for a chip at a certain timepoint. The same steps were applied for all sprouting calculations.

Red fluorescent images of spheroid EC recruitment were processed in ImageJ in a similar manner as previously outlined before importing to MATLAB. A white light image consisting of the mRCC spheroid and EC was used to determine the center point of the spheroid. A custom MATLAB script was used to find the centroid coordinates of the spheroid which appeared darker and more circular than any surrounding structures and thus could be recognized by the software. Manual cropping was employed in situations in which the automated script failed to accurately identify the spheroid. The minimum distance from this center point to the closest image border along the horizontal dimension was used to crop the RFP EC fluorescent images resulting in images of EC centered around the spheroid. A median filter (20×20 kernel) size was applied to the thresholded binary fluorescent images followed by hole filling using the imfill function. The processed images were then divided up into 20 equidistant sections with the EC cellular coverage calculated for each section elucidating the relationship between organoid and EC recruitment. To enable a fairer comparison between chips, 20 sections was used in all cases regardless of individual area analysis. Multiple chips as described in the images were then averaged to assess the population response. Plotting and statistical analysis were performed in either MATLAB or Prism GraphPad (version 9.0.0 (86)) with two-sided students t-tests used to assess the null hypothesis that there was no difference between groups, with p values < 0.05 rejecting the null hypothesis.

PSMA stained on the MFC images were processed in ImageJ using a custom build ImageJ script that thresholds the images leaving only the PSMA stain and applies a median filter (10×10 filter size) to remove any artifacts. Processed images were then imported in MATLAB and analyzed via a custom script that cropped the images to the same size, divided them into 20 equidistant sections, and calculated the percentage of PSMA area in each section. The percentage area values were averaged across the three images. The 20 averaged values were passed through MATLAB *polyfit* function to generate a function describing the PSMA expression as a function of the distance from the tumor center. The *polyfit* function was also applied on the 20 percentage PSMA area values for each image separately. The location of tumor centers was measured manually by inspecting images in MATLAB. Prism GraphPad was used to plot the data obtained from the *polyfit* function in MATLAB.

### Immunohistochemical staining, image quantification, and analysis

Formalin fixed paraffin embedded tumor tissues were processed by the MSKCC Molecular Cytology Core Facility. Tumor sections were stained with a primary mouse monoclonal anti-CD31 (Roche #760-4378) antibody. Next, sections were incubated with secondary Alexa Fluor 647 (Thermo Fisher Scientific # B40958) antibody. Cell nuclei were stained with DAPI (Thermo Fisher Scientific #D3571). Staining was done using automated immunohistochemical staining processor Discovery XT (Ventana Medical Systems). Images were scanned and viewed with 3DHISTECH Ltd. Case Viewer.

CD31 stained images (n=9 control group, n=9 treated group) were processed in ImageJ using a custom build ImageJ script that thresholds the images leaving only the CD31 stain and applies a median filter (10 pixels radius) to remove any noise in the thresholded images. Processed images were then analyzed in MATLAB using a custom script that calculates the percentage of image area occupied by CD31 stain. CD31 stained images from control and treated mice groups were averaged, statistically analyzed, and plotted in Prism GraphPad with the two-sample t-test used to assess the null hypothesis that there was no difference between the two groups, with p values <0.05 rejecting the null hypothesis.

### Assessment of angiogenesis-related proteins on the microfluidic chip

Angiogenesis proteomics was evaluated using an array (R&D Systems #ARY007) to identify angiogenesis-related proteins in culture medium perfusate from the MFC. Medium was collected from the MFC maintained in culture for 15 days in different conditions: EC control perfused with regular medium, EC control co-cultured with mRCC perfused with regular medium, EC control perfused with enriched medium, EC PSMA perfused with enriched medium. Antibodies against 55 proteins were spotted in duplicate on nitrocellulose membranes. 25 mL of culture medium perfusate from each respective condition was diluted and mixed with 25 μL of antibodies detection cocktail. Membranes were incubated for 48h at 4°C with antibodies. Next, membranes were washed to remove unbound material, and stained with IRDye 800CW Streptavidin (LI-COR # 926-32230) at final concentration of 1.0 μL/mL. The Odyssey imaging system was the used to detect differences in expression levels between proteins. Images were manually quantified in ImageJ using identical ROI sizes across all chips with background subtraction performed. Graphing and statistical assessment of the data was performed in Prism, see Supplemental Figure 11. Statistical significance between all conditions was determined using an Ordinary ANOVA test without matching or pairing, comparing the mean of each group via a Tukey statistical hypothesis testing. A family-wise alpha threshold of 0.05 was required for significance with a 95% confidence interval.

### Animal models and vascular targeted therapy *in vivo*

Patient-derived mRCC tumors were implanted into 4-5 weeks old female NOD.Cg-Prkdc^scid^/J mice (Jackson Laboratory) by subcutaneous injection into the upper region of the thigh of 5,000,000 cells in 100 μL of Matrigel (Corning # 356231). Tumor growth was monitored, and mice were imaged with RSOM. To track changers in the vasculature RSOM imaging was performed 24h before mice received the first dose of treatment, and 24h after the last does of treatment. Mice were administered 6 doses of bevacizumab 20 mg/kg (*47*) in 50 μL saline solution or 50 μL of saline, injected every second day for two weeks. RSOM imaging was performed under 2% v/v isoflurane inhalation anesthesia (Baxter #NDC 10019-360-60). Mice were shaved with depilatory cream before (Hair Removal Lotion, Nair) imaging to prevent light absorption and reduced image quality. All animal procedures were approved by the Institutional Animal Care and Use Committee (IACUC) and followed institutional and NIH guidelines.

### Automated quantification of tumor vascular images obtained via RSOM

RSOM images were acquired using the RSOM P50 (iThera Medical, Munich, Germany) at a single wavelength of 532 nm (hemoglobin isosbestic point). Xenografted mice were anesthetized via induction with isoflurane (3% v/v) followed by maintenance at 2% v/v. Centrifuged ultrasound gel dissolved in distilled water (30%) was applied to the imaging area using a wet cotton tipped applicator to avoid bubbles. An area of 12×12×4 mm was then imaged to assess the vascular network within the tumor for both treatment and control groups. Reconstructed images were then exported and frequency split into low (11 - 33 MHz, red) and high frequencies (33 - 99 MHz, green) to increase image fidelity. The green and red channels were then combined and exported as an 8-bit png image for automated quantification in ImageJ. A custom batch processing image analysis script was developed for the automated quantification of vascular networks as recorded by the RSOM. Firstly, the user was prompted to provide the directory location to vascular network images which were then imported and converted to 8-bit grayscale images. Once in grayscale formats the images underwent background subtraction, a tubeness filter was applied along with adaptive median filtering and thresholding. The thresholded image was quantified for area fraction before being skeletonized. The skeletonized image was then used to calculate the vascular network length and vascular network features such as number of branches, junctions, endpoints, slabs. The saved branching network data was then used to assess the tortuosity of each branch (branch length over Euclidean distance). Finally, the thresholded image prior to skeletonization was used to determine the distances between branches (nearest neighbor distance). In all cases the automated script employs built in and open access functions within the ImageJ program and exports a variety of images and excel format files including skeleton overlays for inspection and later analysis. Values for area fraction, network length, NND, branching and tortuosity were then compiled in Prism GraphPad for graphing and statistical analysis.

## Acknowledgements

We thank the Abraham Hakimi Lab from MSKCC for providing us a JHRCC62A patient derived metastatic renal cell carcinoma cells. We thank Andrew Cinquina for assistance with engineering cells. We thank Ning Fan and Afsar Barlas from the MSKCC Molecular Cytology Core Facility for assistance with histology and immunohistochemistry.

## Author information

M.S. designed and performed all experiments, processed, and analyzed the data, and wrote the manuscript. B.M.L., J.C.D. and A.V.D. analyzed confocal microscopy and RSOM images using via scripts in both MATLAB and ImageJ based on established methods and available packages. N.B.P. and J.D. performed selected MFC experiments. A.V. and V.P. engineered cells and preformed FACS. A.O. and C.L.M. provided conceptual input. J.G. supervised the study, provided input for all the experiments and the study concept, and edited the paper.

