## Supplemental Material for "Vascularized tumor on a microfluidic chip to study mechanisms promoting tumor neovascularization and vascular targeted therapies"

**Short title (50 characters):** Microfluidic chip study of tumor vascularization

Magdalena Skubal PhD[1], Benedict Mc Larney PhD[1], Ngan Bao Phung[1,3], Juan Carlos Desmaras[1], Abdul Vehab Dozic[1], Alessia Volpe PhD[1,2], Anuja Ogirala[1], Camila Longo Machado PhD[1,2], Jakob Djibankov[1], Vladimir Ponomarev MD PhD[1,2,3,4], Jan Grimm MD PhD[1,2,3,4,*]

[1] Molecular Pharmacology Program, Memorial Sloan Kettering Cancer Center, New York, NY, USA
[2] Department of Radiology, Memorial Sloan Kettering Cancer Center, New York, NY, USA
[3] Department of Pharmacology, Weill Cornell Medical College, New York, NY, USA
[4] Department of Radiology, Weill Cornell Medical College, New York, NY, USA

keywords: microfluidic chip, neovascularization, targeted therapies, confocal imaging, optoacoustic imaging

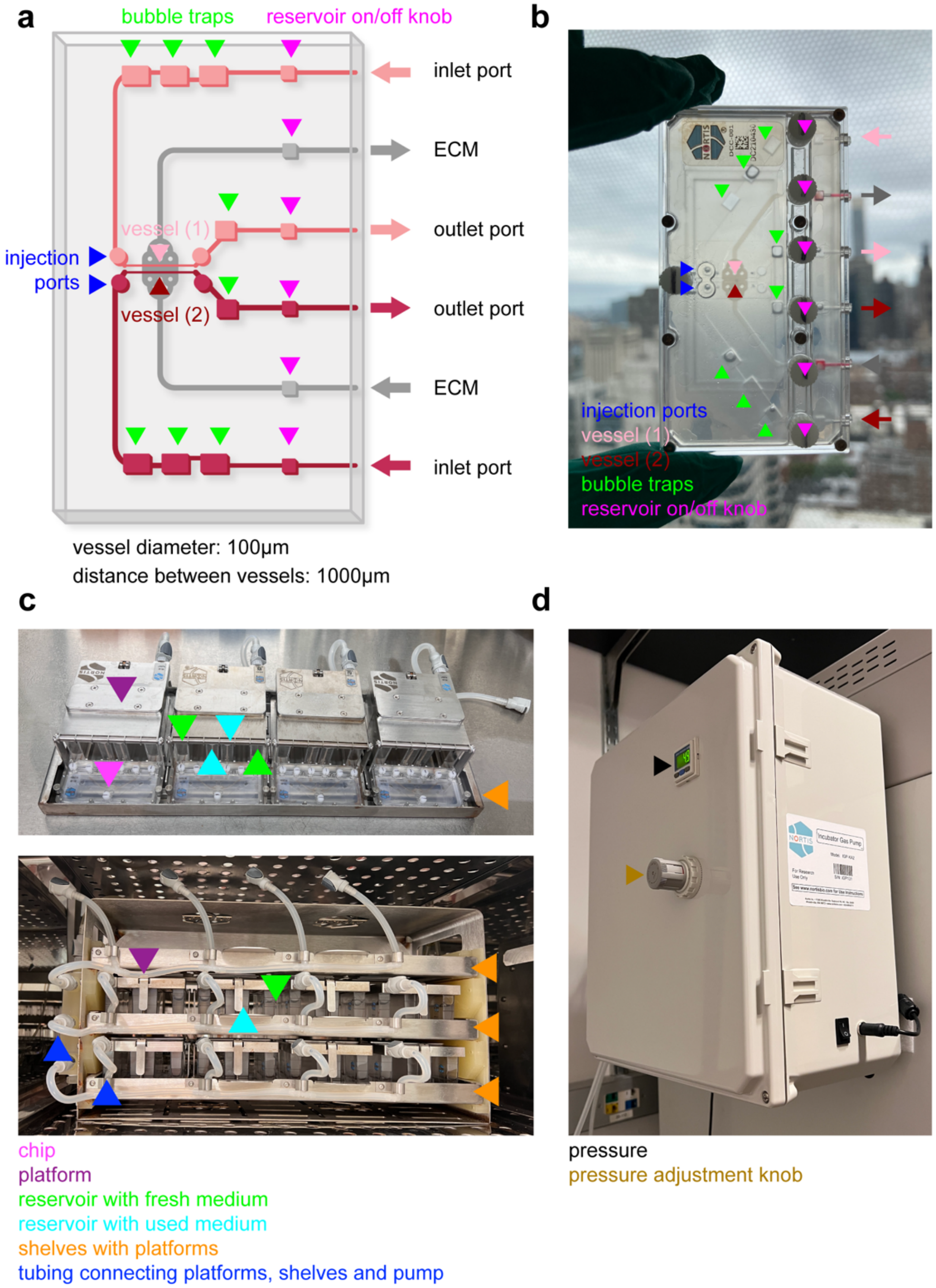


***Supplementary Fig. 1: Double channel microfluidic chip.*** *(a) Schematic visualization and* ***(b)*** *image of double channel MFC built of ECM (extracellular matrix), inlet (fresh medium) and outlet (used medium) ports, vessels, injection ports, bubble traps, reservoir on and off knobs.* ***(c)*** *Single shelf comprised of 4 MFC platforms (top) and image of the MFC system comprised of 3 shelves with 4 chips per shelf within the incubator (bottom). Platform locations, connection tubing, shelving, fresh and used media reservoirs are highlighted.* ***(d)*** *Pumping unit mounted on a dedicated incubator with pressure indicator and pressure adjustment knob.*

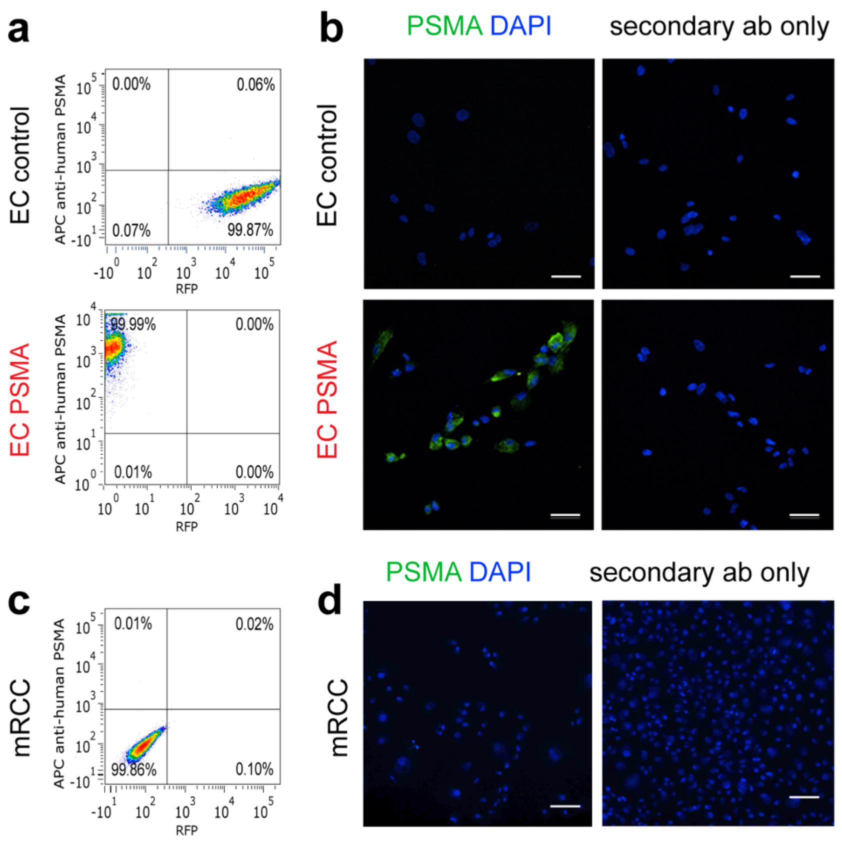


***Supplementary Fig. 2: PSMA expression in endothelial and renal carcinoma cells.*** *(**a**, **c**) Dot-plots from flow cytometric analysis and (**b**, **d**) immunostaining analysis confirming PSMA expression in control EC (EC control), EC engineered by retroviral gene transfer to stably overexpress functional PSMA protein (EC PSMA) and renal carcinoma cells (mRCC). EC control and mRCC are negative for PSMA. Scale bars 100 µm.*

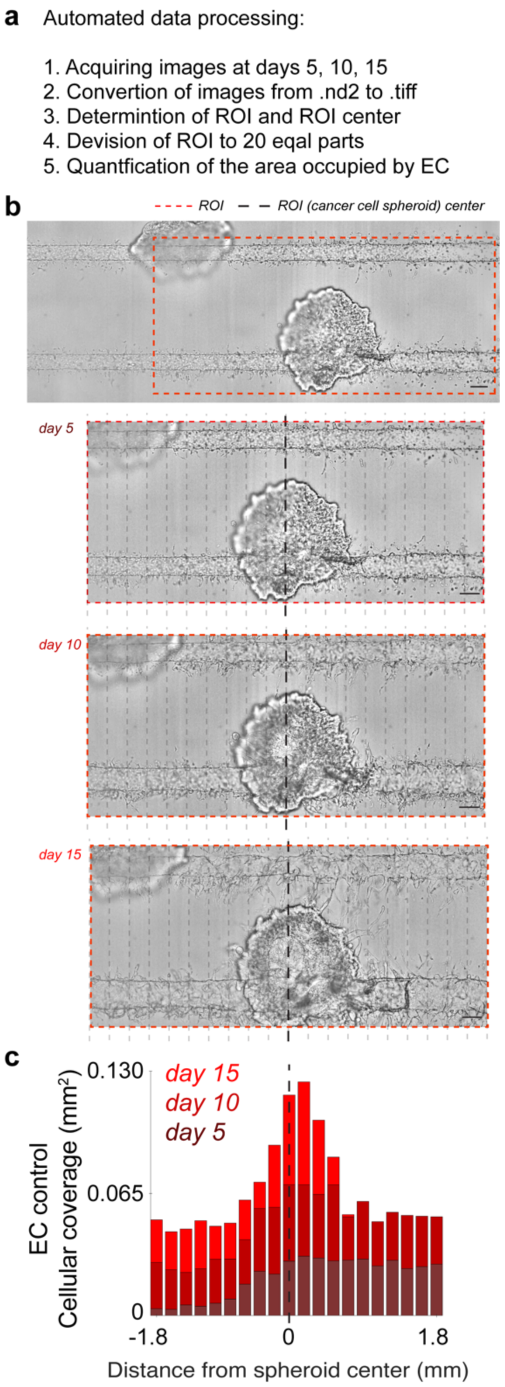


***Supplementary Fig. 3: Quantification of cellular coverage induced by mRCC spheroids. (a)*** *Steps applied to quantify EC coverage: coverage of EC was quantified in the region of interest (ROI) labeled by red dotted lines, the ROI was divided into 20 respective equidistant sections in all cases labeled by gray dotted lines, the ROI center is labeled by black dotted lines and the area occupied by sprouting EC was quantified in relation to the ROI center.* ***(b)*** *Representative images of EC and mRCC spheroid co-culture maintained on chip at days 5, 10 and 15. Scale bars 100 µm.* ***(c)*** *Quantification of the area covered by EC sprouting induced by mRCC spheroid.*

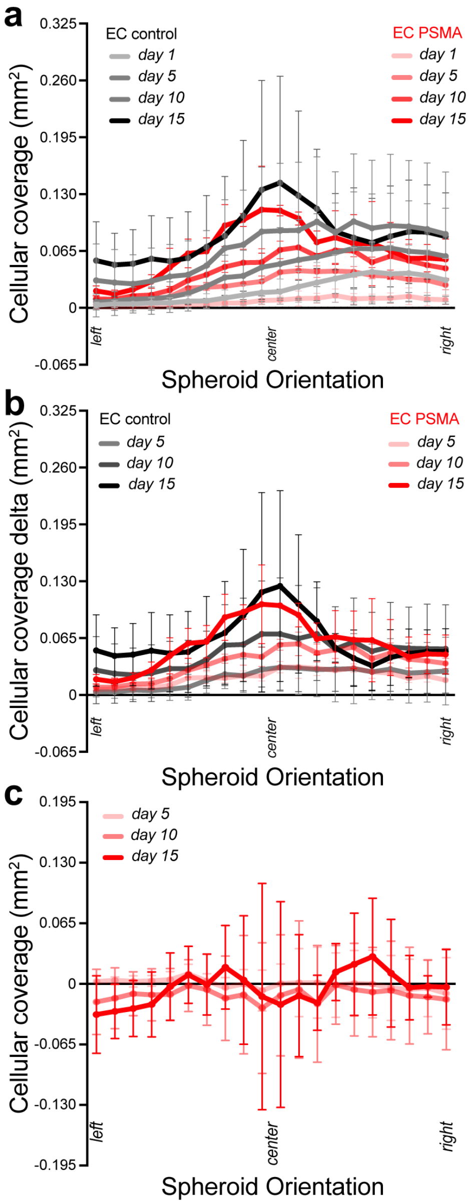


***Supplementary Fig. 4: Cellular coverage of EC control and EC PSMA.*** *(**a**) Plots showing the area covered by EC control and EC PSMA sprouting towards mRCC spheroid at days 1, 5, 10 and 15. (**b**) Plots showing the area occupied by EC control and EC PSMA sprouting towards mRCC at days 5, 10 and 15 after subtraction of day 1 seeding coverage from each respective chip from all other respective timepoints. (**c**) Cellular coverage of EC control coverage values subtracted from EC PSMA values to assess the difference between the groups. No statistically significant regions or timepoints were identified highlighting the EC control similarity to EC PSMA also in line with the induced PSMA expression (see main Figure 2d-e).*

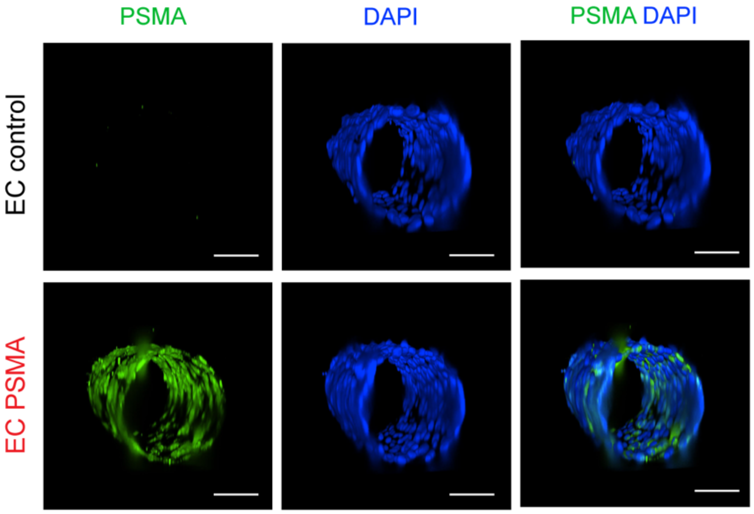


***Supplementary Fig. 5: PSMA expression in EC on microfluidic chip.*** *Immunostaining confirming PSMA expression in EC PSMA (n=1) engineered to stably overexpress functional PSMA protein, and no PSMA expression in EC control (n=1) at day 1. Scale bars 50 µm.*

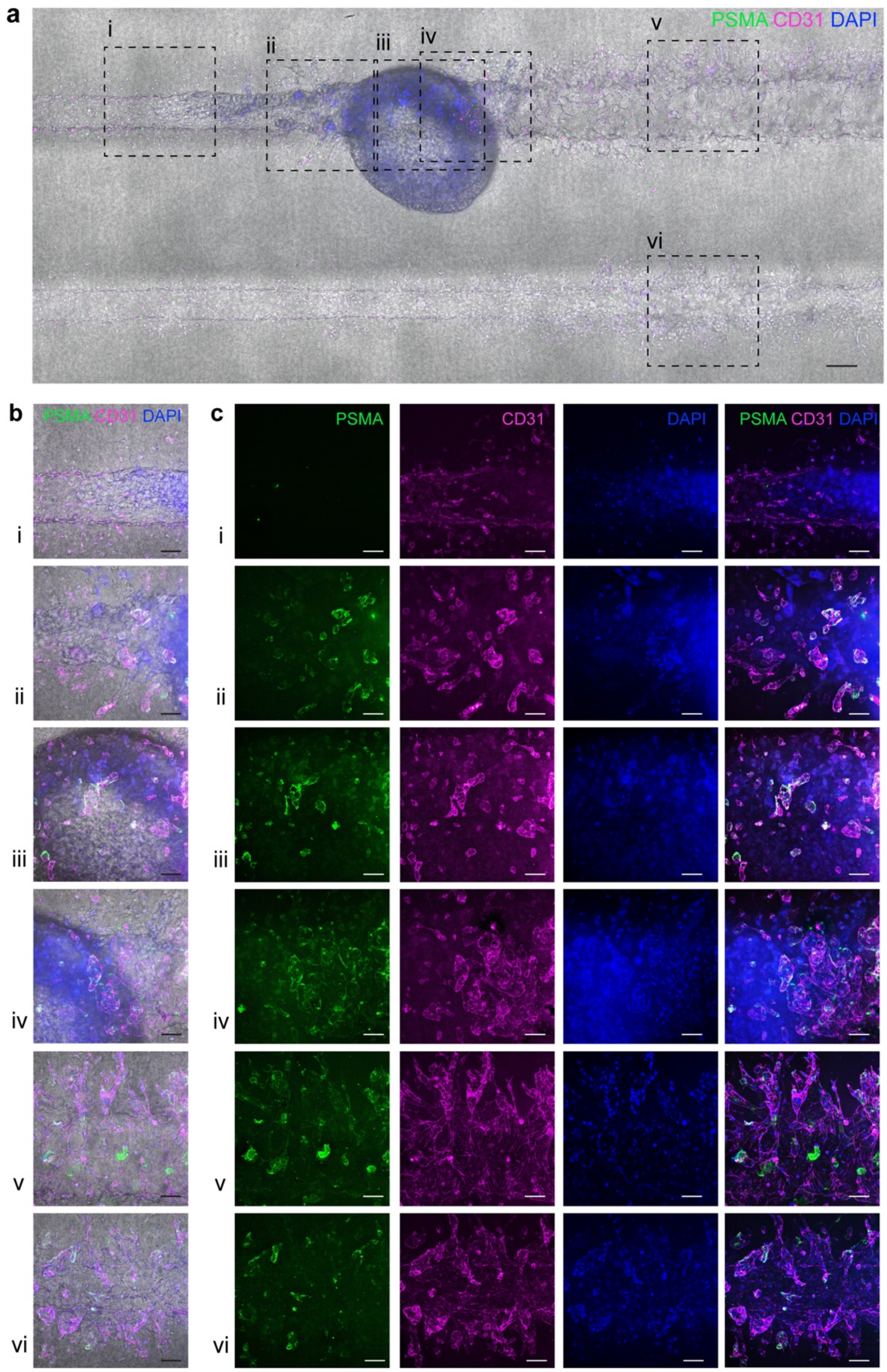


***Supplementary Fig*. 6: Induced PSMA expression on tumor associated neovasculature. (a-c)** *Immunostaining showing PSMA (green) expression induced on control EC (CD31, magenta) co-cultured with mRCC spheroids for 15 days on the MFC. Endothelial and cancer cell nuclei stained with DAPI (blue). Scale bars 100 µm.*

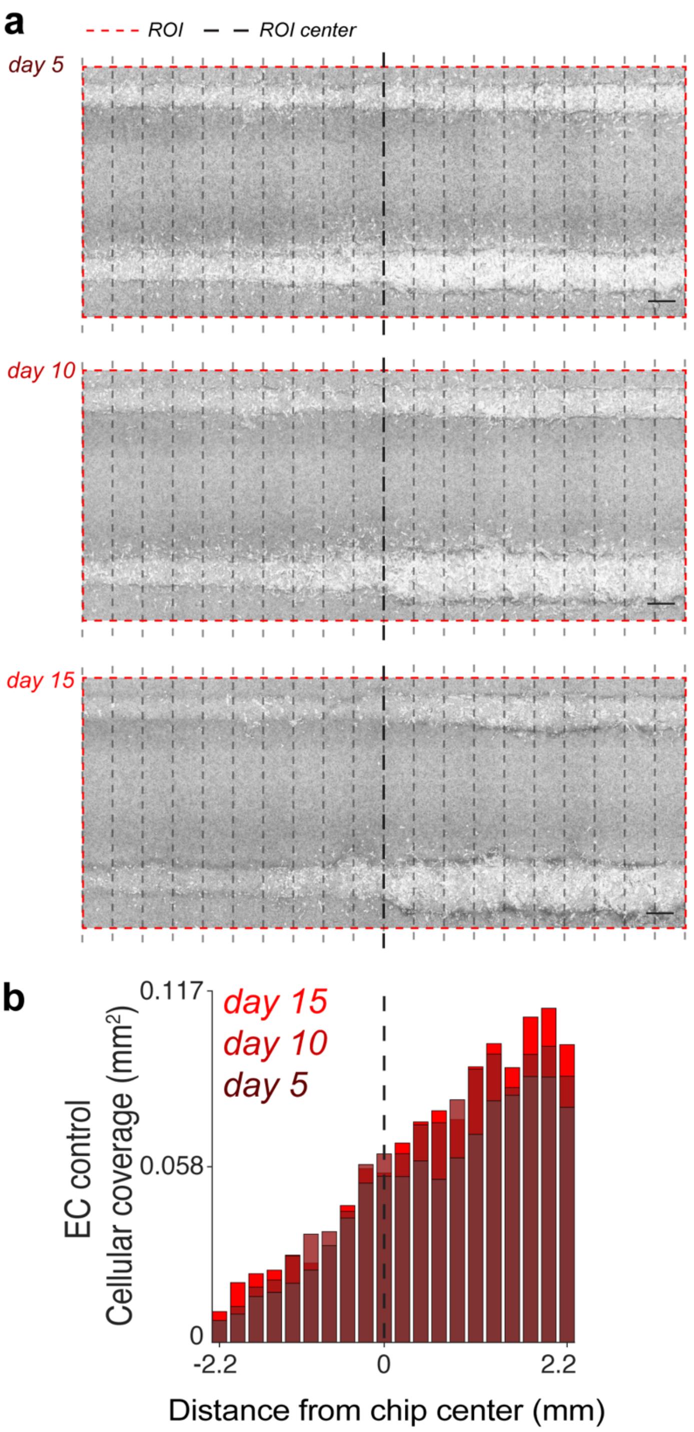


***Supplementary Fig. 7: Quantification of cellular coverage in response to flow effect. (a)*** *Representative images of EC control maintained on chip at days 5, 10 and 15 to visualize flow effects. Scale bars 100 µm. (**b**) Quantification of the area covered by EC sprouting in response to flow effect (n=3). Coverage of EC was quantified in the same manner as in Supplementary figure 3a.*

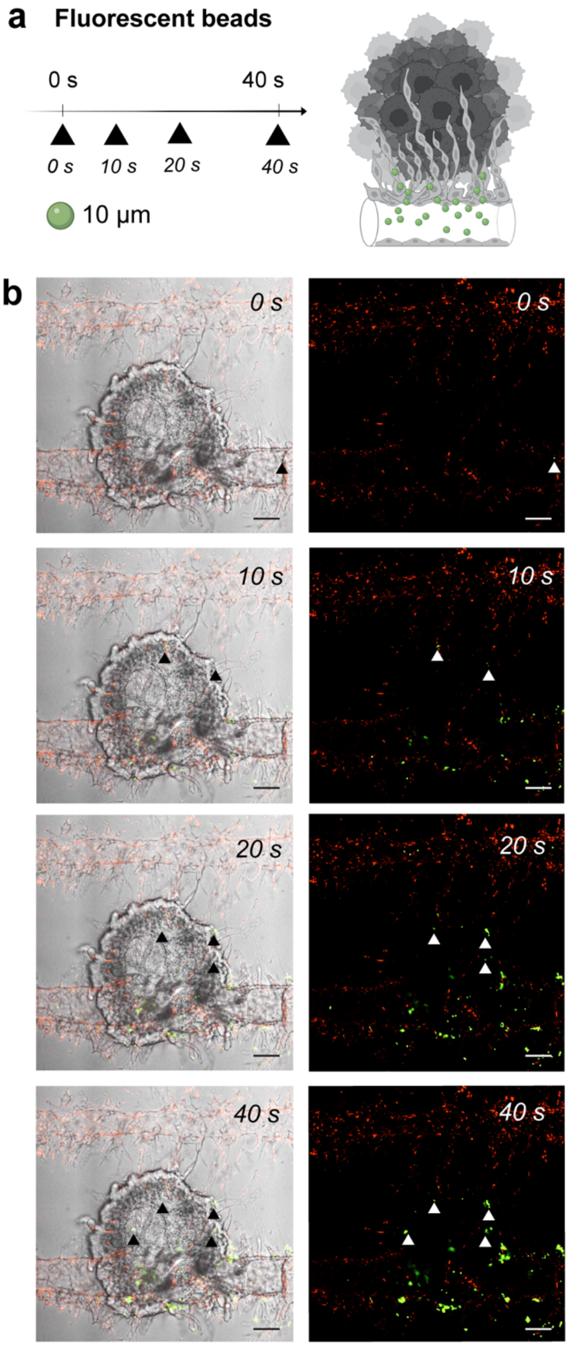


***Supplementary Fig. 8: Mimicking blood flow on the microfluidic chip.*** *(**a**) Functionality of tumor associated blood vessels was tested by perfusion of ø10 µm green, fluorescent beads through tumor associated blood vessels. (**b**) Beads injected into tumor on chip (n=1) with well-developed vasculature (day 15) through the outlet port, entered chip channels and newly formed vessels surrounding the surface of the mRCC spheroid (right panel: beads with EC surrounding mRCC spheroid, left panel: beads with EC) at 0s, 10s, 20s and 40s. Scale bars 100 µm.*

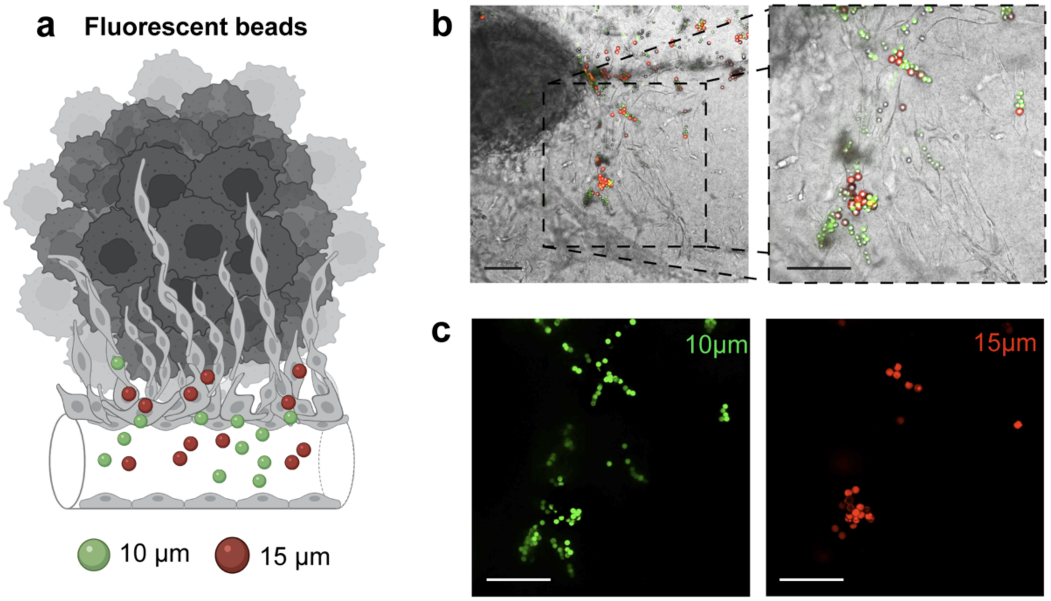


***Supplementary Fig. 9: Mimicking blood flow on the microfluidic chip.*** *(**a**) Functionality of tumor associated blood vessels was tested by perfusion of ø10 µm (green) and ø15 µm (red) fluorescent beads representing sizes in immune cells. Beads injected into tumor on chip (n=1) with well-developed vasculature (day 15) through the outlet port, entered chip channels and newly formed vessels surrounding the surface of the mRCC spheroid. (**b**, **c**) Confocal microscopy images of 10 µm and 15 µm beads traveling through the chip. Scale bars 100 µm.*

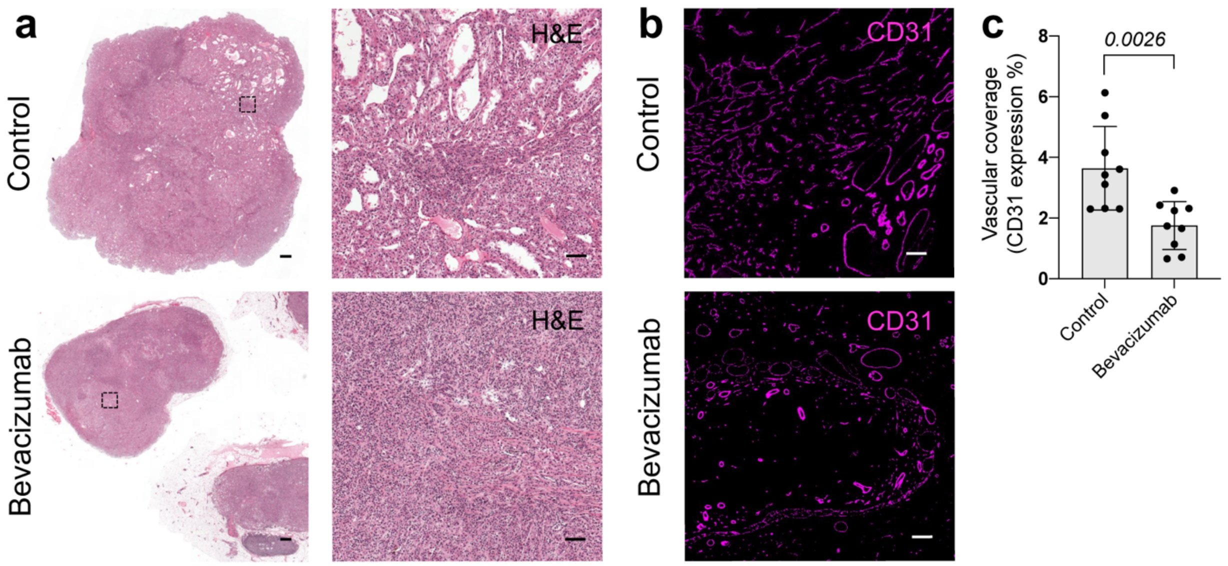


***Supplementary Fig. 10: Vascular targeted therapy in vivo.*** *Immunostaining showing responses to vascular targeted therapy in mRCC tumor model evaluated by treatment with bevacizumab. (**a**) Representative hematoxylin and eosin staining of control and bevacizumab treated tumors. Scale bars 1000 µm and 100 µm, respectively. (**b**) Representative images of CD31 staining of control and bevacizumab treated tumors. Scale bars 100 µm. (**c**) Quantification of vascular coverage after bevacizumab treatment.*

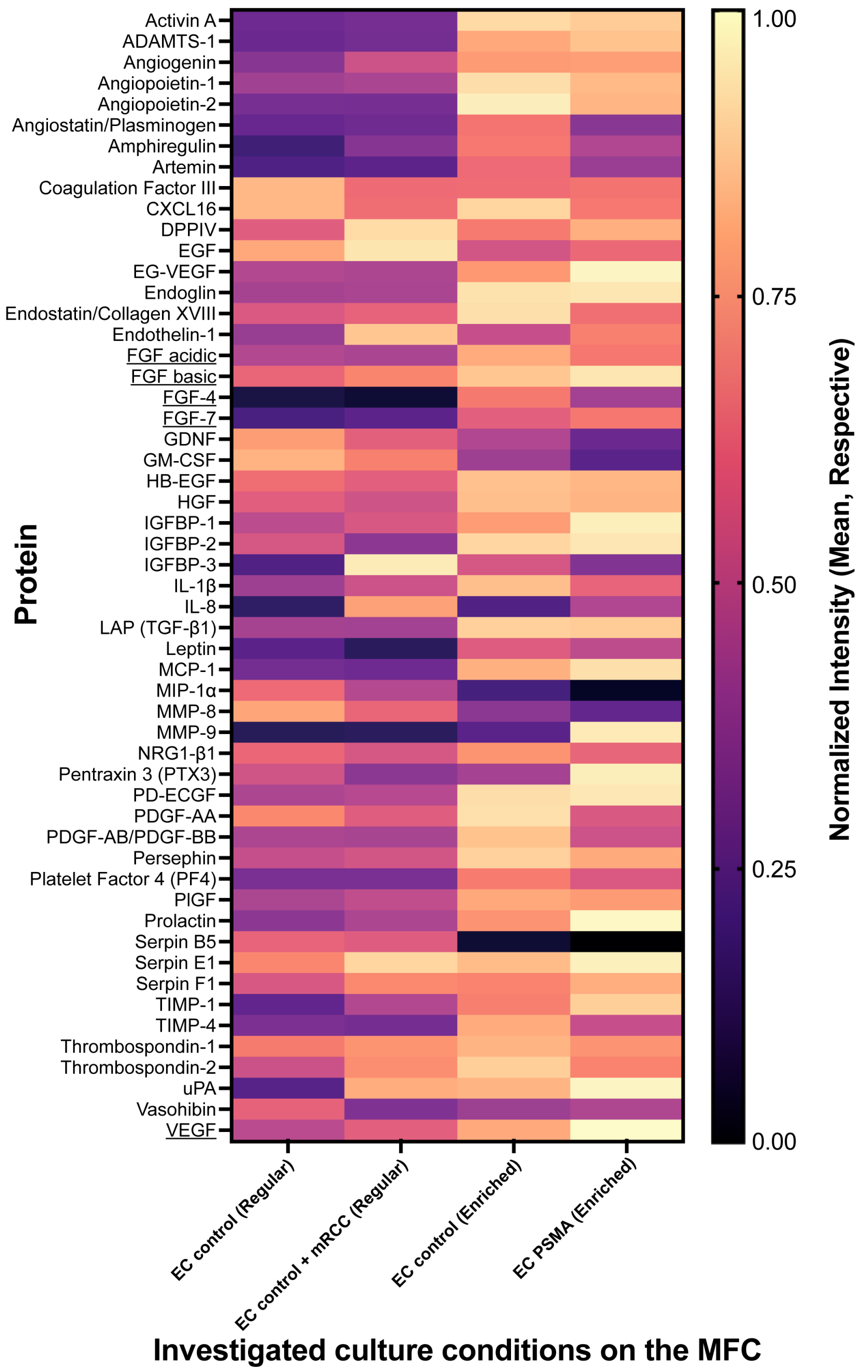


***Supplementary Fig. 11: Microfluidic system and assessment of angiogenesis-related proteins.*** *Heatmap shows the mean values of 55 angiogenesis-related proteins expression in the following conditions: EC control perfused with regular medium (EC control (Regular)) (n=4), EC control with mRCC spheroid perfused with regular medium (EC control + mRCC (Regular) (n=4), EC control perfused with enriched medium (EC control (Enriched)) (n=4), and EC PSMA perfused with enriched medium (EC PSMA (Enriched)) (n=2). Protein expression levels were assessed after 15 days of growth, underlined proteins (note only FGF variations and VEGF) are artificially*

*higher in enriched media. All protein expression has been normalized via division of the highest expression condition for easier visualization of differences. Described significance was only present when $p < 0.05$.*

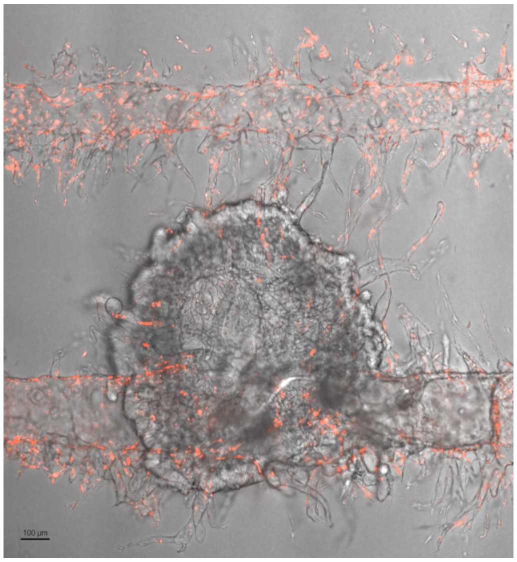


***Video 1: Mimicking blood flow.*** *Fluorescent ø10 µm beads (green) perfused through vascularized tumor on a microfluidic chip at day 15 (EC: red, mRCC spheroid: grey).*

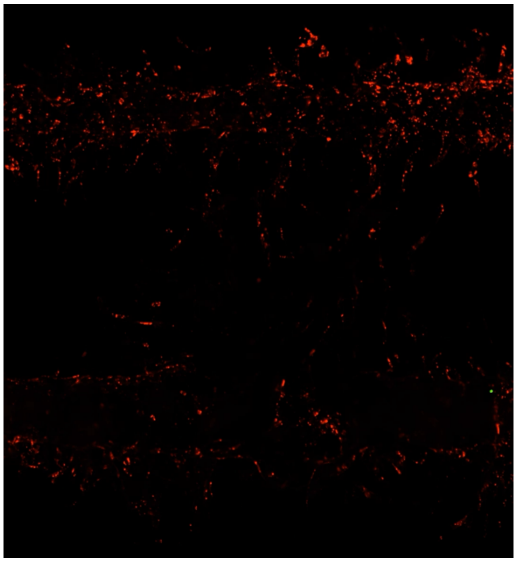

***Video 2: Mimicking blood flow.*** *Fluorescent ø10 µm beads (green) perfused through vascularized tumor on a microfluidic chip at day 15 (EC: red).*

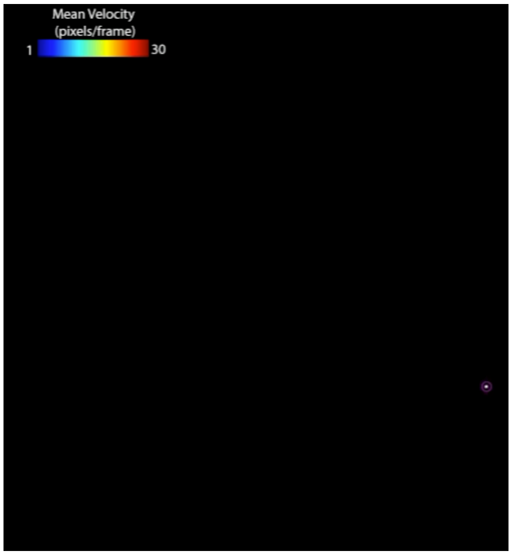


***Video 3: Mimicking blood flow.*** *Tracking of fluorescent ø10 µm beads perfused through vascularized tumor on a microfluidic chip at day 15.*

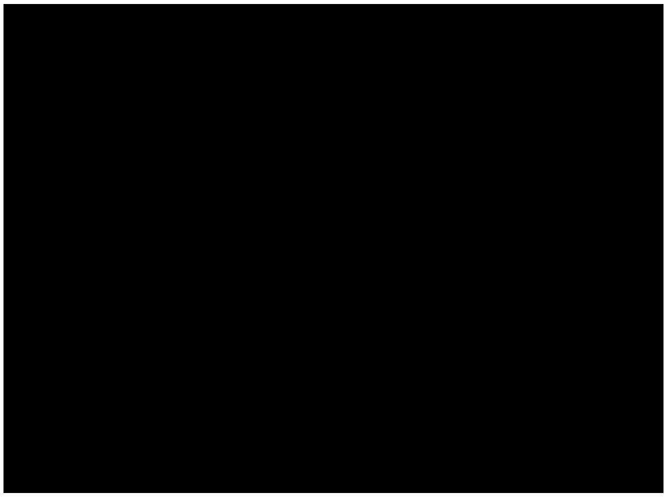

***Video 4: Mimicking drug delivery.*** *Fluorescein (green) perfused (washed in) through vascularized tumor on a microfluidic chip at day 5 (EC: red, mRCC spheroid: grey).*

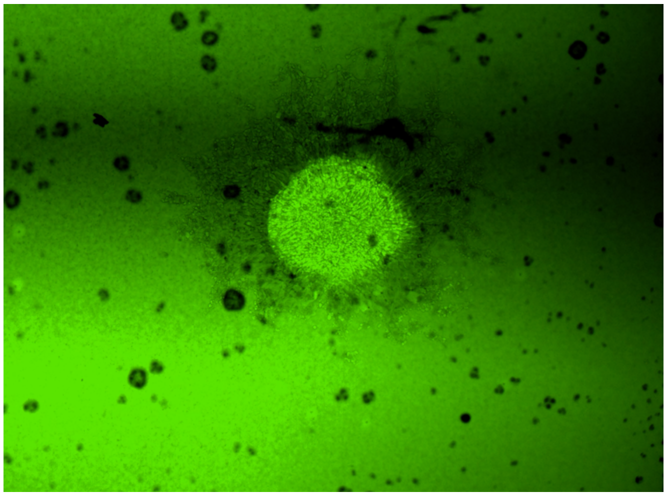

***Video 5: Mimicking drug delivery.*** *Fluorescein (green) perfused (washed out) through vascularized tumor on a microfluidic chip at day 5 (EC: red, mRCC spheroid: grey).*
